# Pitfalls in methods to study colocalization of nanoparticles in mouse macrophage lysosomes

**DOI:** 10.1101/2022.07.21.500664

**Authors:** Aura Maria Moreno-Echeverri, Eva Susnik, Dimitri Vanhecke, Patricia Taladriz-Blanco, Sandor Balog, Alke Petri-Fink, Barbara Rothen-Rutishauser

## Abstract

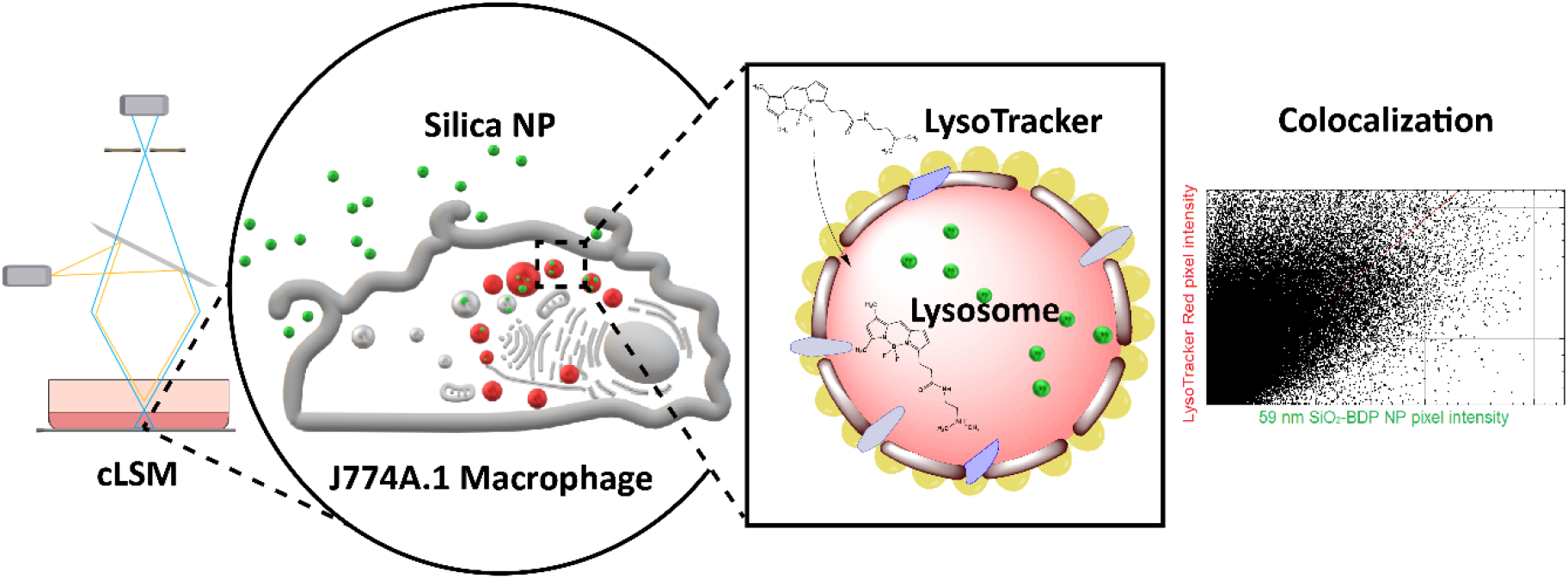

**Background:** In the field of nanoscience there is an increasing interest to follow dynamics of nanoparticles (NP) in cells with an emphasis on endo-lysosomal pathways and long-term NP fate. During our research on this topic, we encountered several pitfalls, which can bias the experimental outcome. We address some of these pitfalls and suggest possible solutions. The accuracy of fluorescence microscopy methods has an important role in obtaining insights into NP interactions with lysosomes at the single cell level including quantification of NP uptake in a specific cell type.

**Methods:** Here we use J774A.1 cells as a model for professional phagocytes. We expose them to fluorescently-labelled amorphous silica NP with different sizes and quantify the colocalization of fluorescently-labelled NP with lysosomes over time. We focus on confocal laser scanning microscopy (CLSM) to obtain 3D spatial information and follow live cell imaging to study NP colocalization with lysosomes.

**Results:** We evaluate different experimental parameters that can bias the colocalization coefficients (i.e, Pearson’s and Manders’), such as the interference of phenol red in the cell culture medium with the fluorescence intensity and image post-processing (effect of spatial resolution, optical slice thickness, pixel saturation and bit depth). Additionally, we determine the correlation coefficients for NP entering the lysosomes under four different experimental set-ups. First, we found out that not only Pearson’s, but also Manders’ correlation coefficient should be considered in lysosome-NP colocalization studies; second, there is a difference in NP colocalization when using NP of different sizes and fluorescence dyes and last, the correlation coefficients might change depending on live-cell and fixed-cell imaging set-up.

**Conclusions:** The results summarize detailed steps and recommendations for the experimental design, staining, sample preparation and imaging to improve the reproducibility of colocalization studies between the NP and lysosomes.

## 1. Background

The use of engineered nanoparticles (NP) for biomedical applications requires a detailed understanding of the NP-cell interaction, NP uptake and intracellular fate. Upon deposition at the outer cell membrane, the NP are internalized via a variety of endocytic mechanisms^1–5^ and pass through an endosomal sorting process. Finally, depending on their properties, NP localize in the subcellular compartments, including lysosomes, endoplasmic reticulum, Golgi apparatus, mitochondria and nucleus^6,7^ or bypass these compartments and end up in the cytoplasm.^8^ Lysosomes are dynamic cellular catabolic organelles responsible for the degradation of macromolecules^9^ and play a key role in the accumulation of internalized materials such as NP.^10,11^ Lysosome dysfunction is associated with multiple diseases, such as cancer, inflammatory and neurodegenerative diseases, and Tay Sachs disease.^12–14^ Therefore, lysosomes are a target for a number of therapeutics^15,16^ and there are several methods available to study the intracellular fate and localization of NP in lysosomes.^17^ Most recognized methods are transmission electron microscopy,^18^ fluorescence microscopy i.e., conventional bright field,^19^ confocal,^20^ and super-resolution microscopy,^21^ correlative microscopy,^22^ focused ion beam-scanning electron microscopy (FIB-SEM)^23^ and lysosomal isolation.^24^ The choice of each method strongly depends on NP physicochemical properties (i.e., core material, size, shape, surface charge, porosity, surface roughness, surface area, crystalline structure, etc.)^25^ and complexity of the intracellular environment.^26^ In this study, we focused on confocal laser scanning microscopy (CLSM), due to the possibility of imaging fluorescently labelled NP, fast sample preparation, obtaining 3D spatial information without the need for physical sectioning of the sample, live cell imaging, and the possibility to obtain information about NP colocalization with cellular compartments.

Different approaches are available to visualize lysosomes within live and fixed cells.^27^ The most used are the LysoTracker probes, Lysosomal-Associated Membrane Protein (LAMP) antibodies^28^ and cells transfected with fluorescent constructs targeting sequence for LAMP.^29^ Here, the focus is on the LysoTracker probes due to the possibility of dynamic lysosome imaging in unfixed cells, low phototoxicity, and ease of use for long term imaging. Considering its many advantages, this approach is widely used staining in research.^28,30–33^ LysoTracker probes contain a conjugated multi-pyrrole ring structure with a weakly basic amine, which gets protonated and accumulates in the acidic environment of the lysosomes and late endosomes.^34^ LysoTracker probes has some drawbacks: i) Low pH is not unique as late endosomes can have a similar pH as lysosomes (∼ 4.5 - 6.5) end, ii) Fluorophore instability^34^, e.g. LysoTracker Red can undergo photoconversion to a green-emitting fluorophore,^34,35^ iii) Variability in intracellular NP trafficking between individual cells (the time for NP to internalize into lysosomes can range from 15 min to several hours),^20^ **iv)** LysoTracker probes and LAMP signal miscorrelation may occur in some cell lines.^28^

Given these challenges, further experimental and observation-based biases must be avoided to provide more reliable data interpretation. Moreover, there are other experimental factors that can lead to misinterpretation of biological results, particularly those that rely on fluorescence and image analysis of live or fixed cells.

Colocalization analysis is a common approach to evaluate NP trafficking and accumulation in lysosomes. Based on optical sections recorded by CLSM, colocalization analysis requires statistical methods.^36^ The data consists of two channels, such as i) the lysosome visualized by LysoTracker and/or LAMP and ii) the fluorescently labelled NP. Fluorescence intensity data is obtained from each recorded pixel between the LysoTracker/LAMP channel and fluorescently labelled NP. Such analysis provides information about co-occurrence (i.e., the extent of spatial overlap between the two fluorophores), as assessed by Manders’ coefficients^37^ and cross-correlation (the functional relationship between the signal intensities of the two fluorophores), as described by Pearson’s correlation coefficient (PCC).^38^ Additional measures and tests may include the Spearmann’s Rank Coefficient^39^, Costes’ randomization approach^40^, Intensity Correlation Quotient^41^ or various object-based measures.^42^ For accurate data interpretation, a combination of multiple metrics is recommended. However, such sound interpretations are only achievable if biases in data gathering are avoided. In our microscopic investigations on intracellular trafficking of NP, in particular to the lysosomes, we encountered several methodical and technical pitfalls when using probes to label lysosomes, each leading to misinterpretation and bias of the lysosomal labelling. Here, we describe the most common pitfalls and provide recommendations on how to circumvent them to yield accurate conclusions.

## 2. Results and discussion

The purpose of the present study is to determine the necessary steps for reliable colocalization of NP within lysosomes with a focus on CLSM. J774A.1 mouse macrophages were used as a model cell line for professional phagocytes, since they are capable of phagocytosis (uptake of particles larger than 1 μm)^43–45^ but are also a well-known model for endocytosis research.^30,46^ In that sense this is an ideal cell line to study a wide spectrum of particle types suited for the purposes of this project. Cells were exposed to amorphous silica NP because they can be synthesized in different sizes and labelled with a variety of different fluorophores, allowing their tracking in living cells.

### 2.1. Fluorescent lysosomal probe characteristics

When designing an experiment, the key experimental parameters (i.e. LysoTracker probe selection and the stability of fluorophores as well as equipment available) should be taken into consideration. Since lysosomes were visualized by labeling the cells with LysoTracker probes, we first extracted the main characteristics of LysoTracker probes **(Table 1)**. Experiments must be designed in a way to minimize fluorescence cross-talk (i.e., overlap of fluorochromes’ excitation spectra) and bleed-through (i.e., overlap of fluorochromes’ emission spectra), either by using a free online spectral analyzer (available by Thermo Fisher Scientific) or experimentally **(Figure S1A)**.^47^

**Table 1.** Main LysoTracker probes characteristics and experimental parameters.

| LysoTracker* | Excitation<br>(nm)<sup>1</sup> | Emission<br>(nm)<sup>1</sup> | Recommended<br>concentration (nM) | Incubation<br>time (37°C) | Ref. |
| --- | --- | --- | --- | --- | --- |
| Green (DND-26) | 504 | 511 | 50 | 15-30 min | 48-50 |
| Blue (DND-22) | 373 | 422 | 50-100 | 30 min – 1.5h | 51-53 |
| Red (DND-99) | 577 | 590 | 10-50 | 15 min – 4h | 20,31,54 |

<sup>1</sup> Wavelength. The information in this table refers to LysoTracker probes purchased from ThermoFisher Scientific.

### 2.2. Influence of phenol red in cell culture medium on fluorescence intensity of LysoTracker probes

Our first attempt was to investigate whether the presence of phenol red, a routinely added pH indicator, in the culture medium influences the fluorescence intensity of the red, green and blue LysoTracker probes. Cells were grown either in complete Roswell Park Memorial Institute medium, supplemented with 10% fetal bovine serum (FBS), 1 % (v/v) L-glutamine and 1 % (v/v) penicillin/streptomycin (cRPMI) with or without phenol red. After 24 h of incubation, the cells were exposed to LysoTracker probes and the effect was monitored under continuous live cell imaging. A representative image of an individual cell for each LysoTracker probe can be found in **Figure S2**. Our results show that the presence of phenol red decreases the relative signal intensity and biases the fluorescent signal as demonstrated by the significant differences in the raw integrated density (i.e., sum of pixel values) of LysoTracker probes **(Figure 1)**. This distorts the correlation coefficients and the interpretation of the colocalization results. The phenol red emits light in the blue and red region of the spectrum, causing mainly interference with the blue and red LysoTracker probes spectra **(Figure S1A)**. On the other hand, no significant impact of phenol red was detected on the intensity of LysoTracker Green. Based on these observations, we suggest using phenol red-free cell culture medium for experiments involving LysoTracker probes. In case the use of phenol red-containing medium cannot be avoided, recommended alternative is to use LysoTracker Green instead.

**Figure 1.**
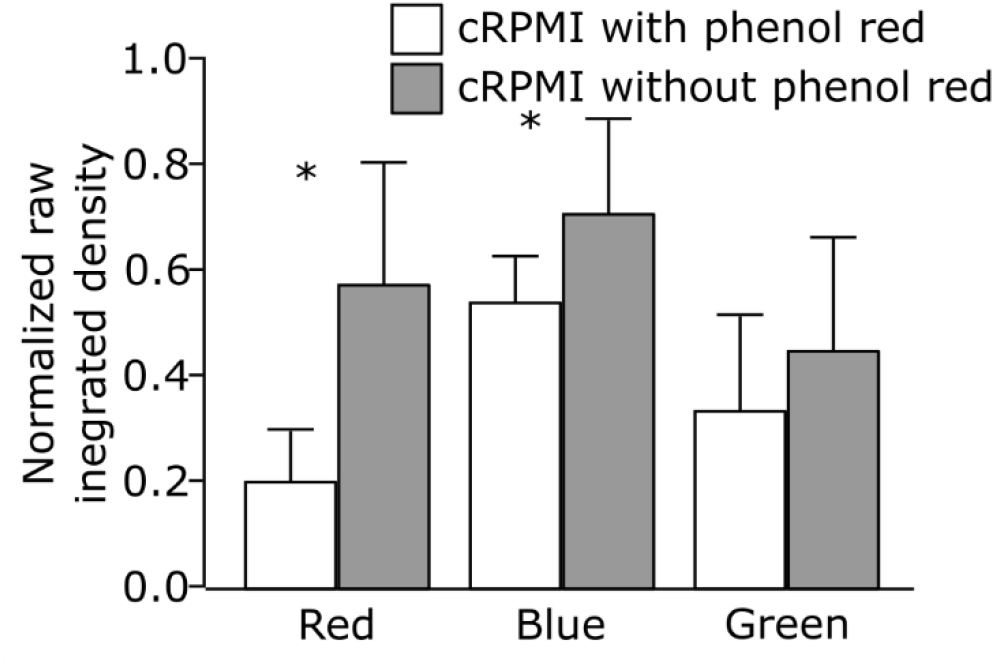
Comparison of the normalized raw integrated densities of three different LysoTracker probes (red, blue and green) obtained by cell imaging in cRPMI with phenol red (white bars) and without phenol red (gray bars). Error bars are standard deviation, n=10 cells per treatment. * p<0.05 by one-way ANOVA.

### 2.3. Stability of LysoTracker Red probe under continuous and staggered imaging set-up

The ability to trace the behaviour of lysosomes over longer periods opens a possibility to study cellular processes upon prolonged and repeated exposure to NP. The suitability of LysoTracker probes for prolonged imaging was evaluated by monitoring the LysoTracker Red fluorescent signal over a period of 24h in a continuous experimental set-up. J774A.1 cells were first stained with LysoTracker Red for 1 hour, the medium was replaced with phenol red-free medium, and cells were imaged live for 24 hours at 20 minutes intervals. Randomized datasets^17^ were recorded.

In a continuous imaging set-up, the fluorescent signal of LysoTracker Red did not significantly alter over the first 8h. Longer periods were accompanied by a decrease in the raw integrated intensity of the LysoTracker Red and increased cell death over time **(Video S1)**. This observation is supported by Chen et al., who demonstrated that prolonged intracellular accumulation of LysoTracker probes causes an increase in intracellular pH and enhances quenching of the fluorescent dye.^27^ However, as assessed by lactate dehydrogenase (LDH) assay **(Figure 2A)** we did not observe any significant changes in LDH release in the medium from LysoTracker treated cells compared to the control cells. Hence, adding LysoTracker probe does not cause cytotoxicity within the time range investigated. Therefore, we conclude that the cytotoxicity is rather influenced by phototoxicity, an effect caused by extensive imaging.

**Figure 2.**
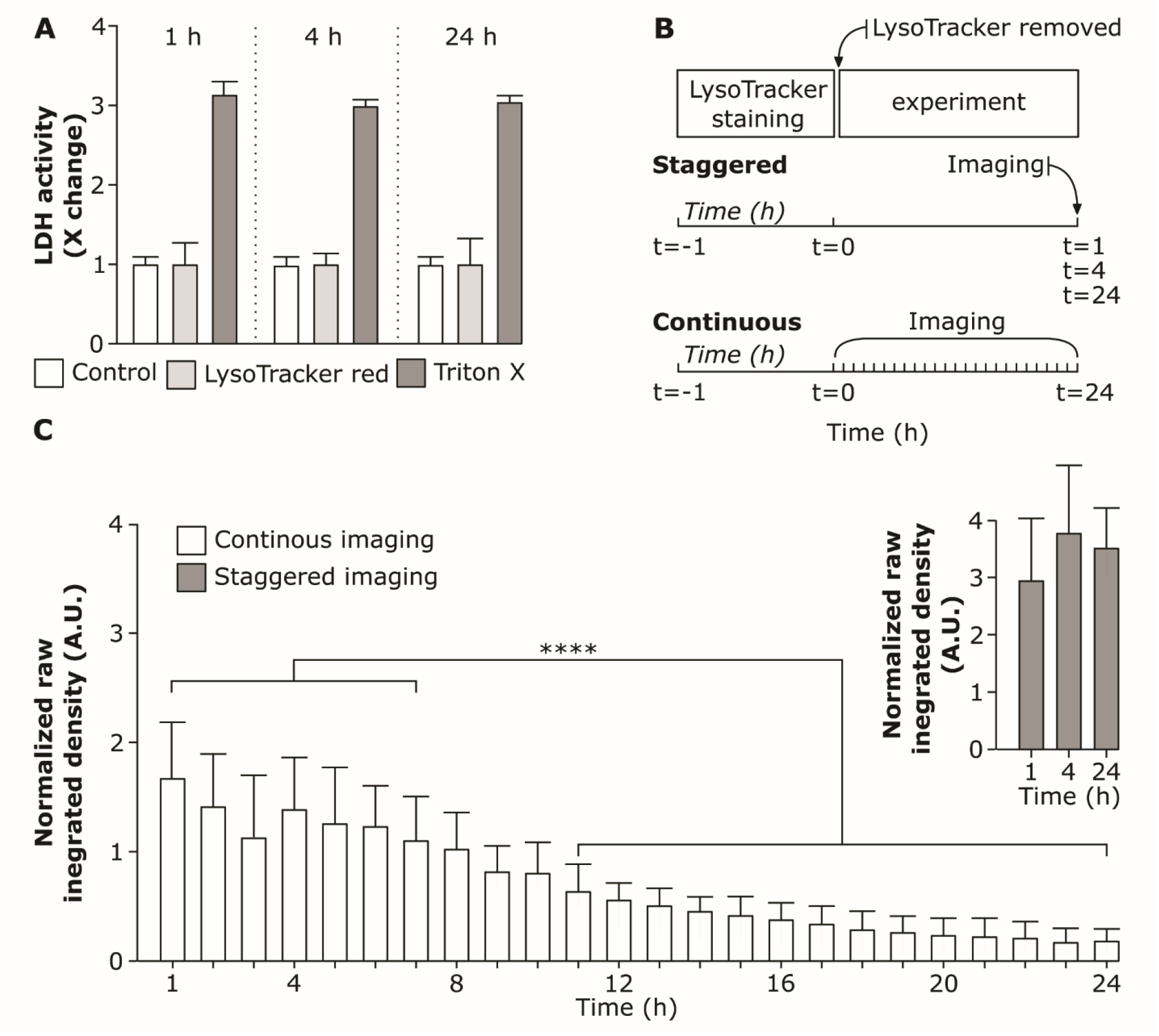
Stability of LysoTracker Red probe under different imaging conditions. **A**. Cell viability after incubation with LysoTracker Red probe for 1 h, 4 h and 24 h assessed via membrane rupture assay (lactate dehydrogenase; LDH) and presented as fold change over untreated cells (control). Data shows the mean of three replicates. Cells exposed to 0.2% Triton X-100 (v/v) served as positive control for LDH assay. **B**. Schematic representation of staggered experimental set-up (cells were stained for 1 h with LysoTracker Red probe. The supernatant containing LysoTracker Red was removed and cells were further incubated and imaged at 1 h, 4 h and 24 h as hyperstacks) and continuous set-up (cells were stained for 1 h with LysoTracker Red probe and imaged continuously for 24 h as hyperstacks). **C**. Comparison of raw integrated densities of LysoTracker Red probe versus time for continuous and staggered imaging. Each time-point (hour) shows the raw integrated density measured for each cell individually. The whiskers are the standard deviations. **\*\*\*\*** (P < 0.05) is the significant difference analyzed by One-way ANOVA.

To test whether there is a difference in LysoTracker Red signal intensity depending on the imaging conditions an additional staggered experimental set-up was conducted as shown in **Figure 2B**. We observed that the raw integrated density does not differ significantly among the staggered imaging set-up of 1h, 4h and 24h after LysoTracker probe incubation **(Figure 2C)**. However, a significant decrease in raw integrated density of LysoTracker Red signal was found in continuous live-cell imaging set-up and the effect accumulated with increasing imaging time. This suggests that the fluorescent intensity of the LysoTracker Red probe drops mostly because of the photobleaching effect due to continuous imaging for a prolonged time.^55^ The effect might be cell-specific and is affected by the imaging settings.

### 2.4. Physicochemical characteristics of silica nanoparticles

Once the LysoTracker probe stability and imaging set-up is optimized, the selection of the optimal NP to assess the specific research question is important. For our experiments, we selected synthetic amorphous silica NP of different sizes and labelled with different fluorophores.

**Table 2** represents the main physicochemical characteristics of different sizes of silica (SiO_2_) NP (59 nm, 119 nm and 920 nm) used in this study; NP of 59 nm and 119 nm size range are internalized into cells mainly via pinocytosis while 920 nm SiO_2_ particles are taken up by phagocytosis.^56^ A concentration of 20 μg/mL of the NP suspension was chosen as this concentration does not impair cell viability^32^ and all the particles used in our studies were negative for the endotoxin test (data not shown). Comparison of the emission and absorption spectra of the different (nano)particles is shown in **Figures S1B and S1C**.

**Table 2.**
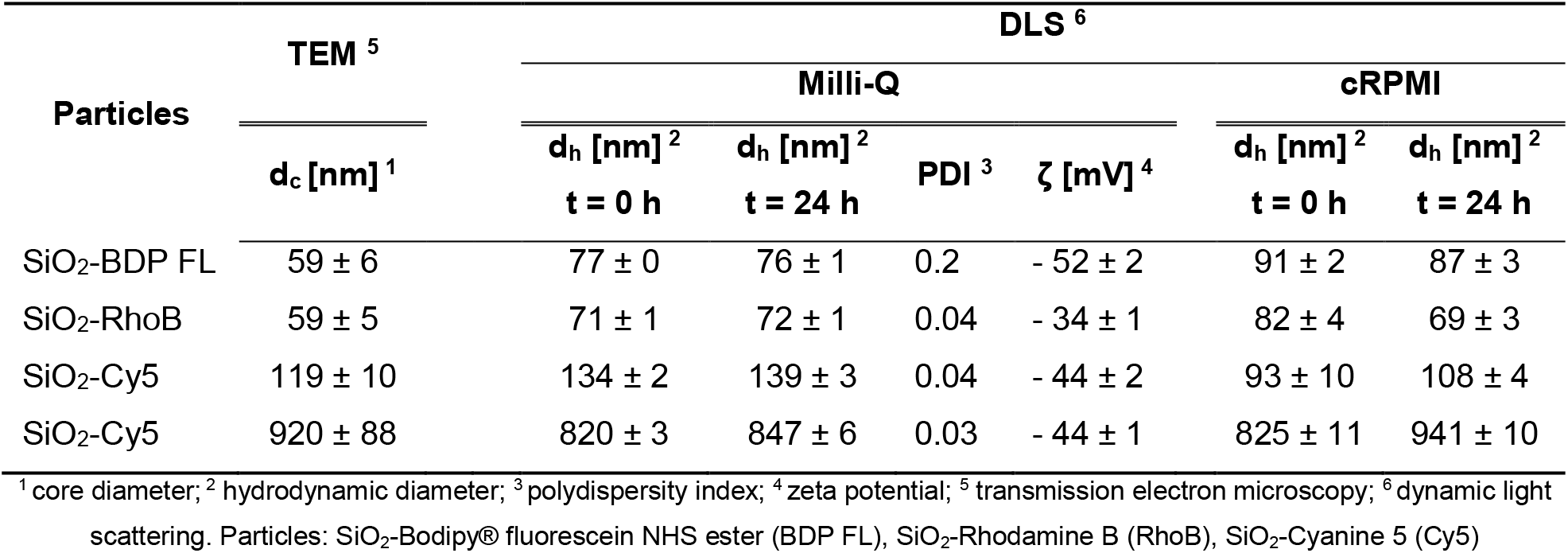
Representation of the main physicochemical characteristics of SiO_2_ particles.

The core particle diameter was measured by transmission electron microscopy **(Figure S3)**, whereas the hydrodynamic diameter was assessed both in Milli-Q water and complete medium via dynamic light scattering (DLS). The corresponding histograms representing particle size distributions are shown in **Figure S4A and S4B**. All SiO_2_ particles were colloidally stable in Milli-Q and in cRPMI over 24h of incubation time.

### 2.5. Confocal microscopy imaging and post-processing

Even optimally designed experiments – no cross talk, no bleed through – and sample preparation can be undone by errors during sample imaging.^57^ Small variations in the settings can significantly alter the correlation coefficients.^58^ Such effects are shown in **Figure 3**. Using an optimally imaged, real dataset, these biases were demonstrated by simulating a set of different imaging conditions for the following data aspects: spatial resolution, slice thickness, pixel saturation, and dynamic range **(Figure 3)**.

**Figure 3.**
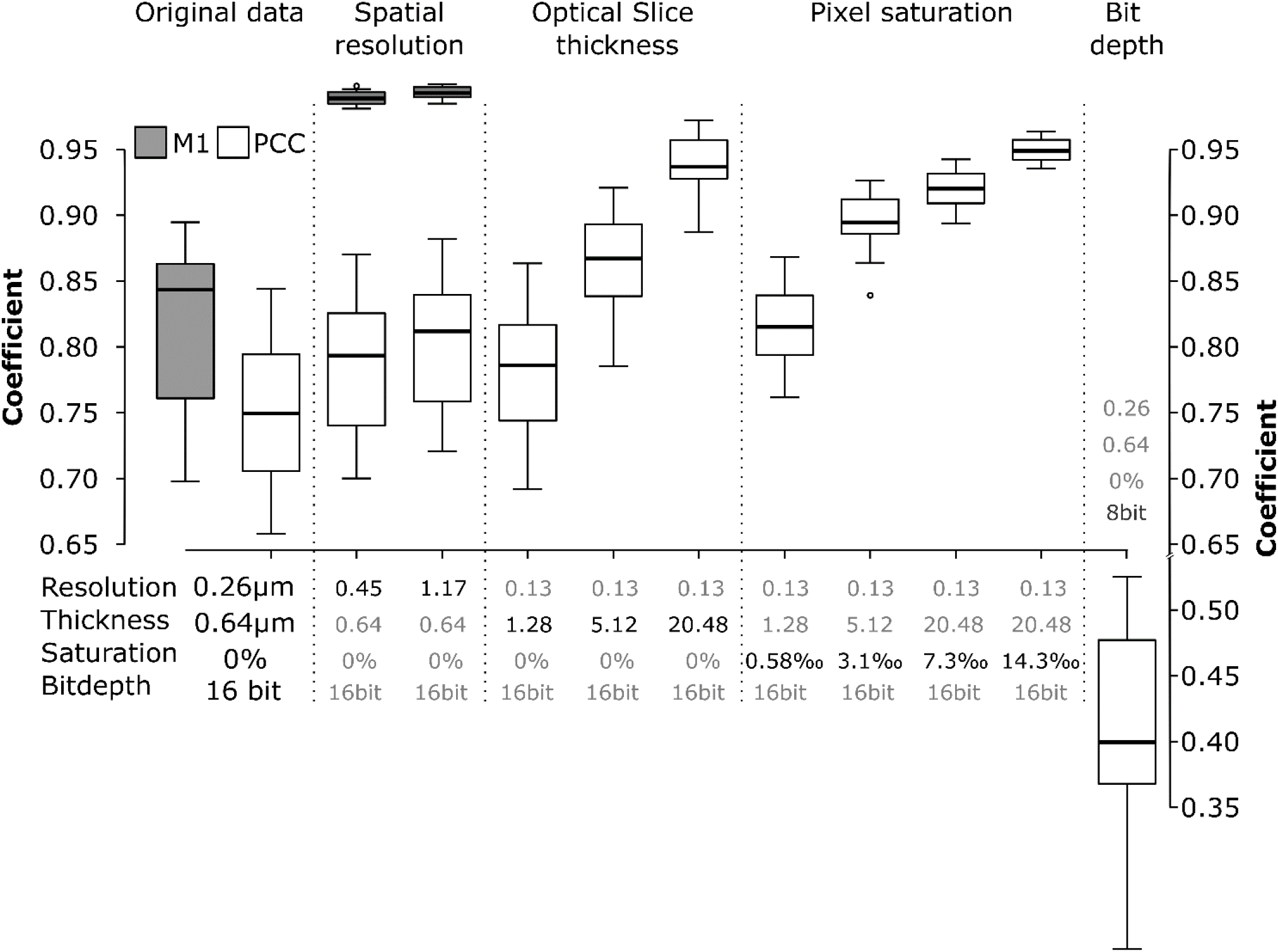
Effect of spatial resolution, optical slice thickness, pixel saturation and bit depth on the Pearson correlation coefficients (PCC, white boxes) and Manders’ M1 coefficient (gray boxes). The data correspond to the experimental set-up where cells were exposed to 59 nm SiO_2_-BDP FL NP for 13 h, fixed and stained with Alexa Fluor 647-conjugated LAMP-2 antibody. The error bars represent the standard deviations. All the images were analysed using ImageJ with the JACoP plugin comparing the correlation coefficients after each specific simulation.

Lowering the resolution by decreasing XY-pixel size (i.e., the distance between points on a specimen, also known as “downsampling”) caused a decrease in Pearson’s correlation coefficient (PCC) and especially in overlap (Manders, M1). Lower resolutions cause the integration of fluorescence signals and thereby affect the interactions between the two channels. This is most clear in the overlap (M1), shown in the grey boxplots of **Figure 3 / Spatial resolution**. Therefore, the optimal resolution to image the objects must be taken into consideration when setting up the experiment.

Slice thickness of the optical slices also affected the colocalization parameters. Simulation of the opening of the pinhole - causing lower axial resolution, i.e. Z-axis thickness - yielded an increase in colocalization measures as objects along the Z-axis are increasingly projected on top of each other and end up falsely overlapping **(Figure 3 / Optical slice thickness)**.

A brighter illumination will improve the signal to noise ratio of the fluorescence signal, which may yield a lower variance of the correlation coefficients. However, the saturation of pixels will bias the result.^59^ Saturation occurs when the fluorescence intensity exceeds the acquisition or storage limit of the detector. Such clipping of data has a significant impact on correlation routines, such as PCC, which try to explain the intensity of the first channel with the information of the second. Our simulations show that even a fraction of saturated pixels as low as 0.3 % of the total pixel number will significantly alter the colocalization **(Figure 3 / Pixel saturation)**. The pitfall of saturation can be completely circumvented by using photon-counting detectors,^60^ which do not convert the signal to binary output (see below) but store the actual number of photons that have arrived at the detector. However, such detectors are relatively new in the field and not yet widespread. Standard GaAsP-PMT detectors used in CLSM convert the analogue fluorescent intensities to a digital signal and store it on a computer. Usually, you will get the choice to save the data with an information depth ranging from 8 bit (2^8^, i.e., 256 different intensity values) up to 16 bit (2^16^, i.e., 65536 different intensity values). Recording data in 16 bit but saving the data at 8 bit, i.e., compressing the information, may cause significant shifts in PCC **(Figure 3 / Bit depth)**.^61^

Before proceeding with further experiments, all microscopy settings, such as resolution, optical slice thickness, saturation and bit depth should be taken into consideration for the reproducibility of the data.

### 2.6. Analysis of nanoparticles within the lysosomes

In this section, we describe four different experiments illustrating the colocalization of silica NP with lysosomes and the corresponding interpretation. The PCC, M1 and M2 coefficients are expressed as a fraction / index of NP colocalized with lysosomes. The experimental design is matched to the specific aim of each study.

#### Experiment I: The effect of continuous live cell imaging on colocalization

Understanding NP localization in lysosomes after their cellular uptake is crucial for their use in biomedical applications.^62^ The advantage of live cell imaging is to get a better insight into intracellular NP trafficking,^20^ and accumulation. The variability in NP uptake dynamics depends on the cell type^43^ and cell division.^20,30,63,64^ The following experimental approach enabled us to investigate whether NP colocalization with lysosomes varies over time under a continuous imaging set-up. J774A.1 cells were initially stained with LysoTracker Red for 1 h and then sequentially exposed to 59 nm SiO_2_-BDP FL NP for a duration of 4h to 8h **(Figure 4A)**. The colocalization was assessed using PCC as an indicator of functional relationship and Manders’ coefficients (M1 and M2) as quantifiers of spatial overlap between fluorophores. PCC values indicate that approximately 0.5 (PCC) of intracellular NP fraction colocalized with lysosomes and there were no significant differences in PCC between 4 h to 8 h **(Figure 4B)**. However, the PCC does not provide detailed information about the distribution of NP arriving into lysosomes and the fraction of remaining empty lysosomes **(Figure 4B)**. In contrast, M1 and M2 coefficients vary between the different time points **(Figure 4C)** and provide information on the fraction of the NP co-occurring with lysosomes within the resolution of a single voxel. We observed that less than 0.1 of NP fraction (M1) reached the lysosomes within the first 4 h of exposure **(Figure 4C)**. The fraction of NP that accumulated in the lysosomes increased gradually over time as confirmed by increasing M1 fraction up to 0.5 after 8 h of incubation. However, there was a variability in the fraction of lysosomes containing NP over time (M2). At 8 h time-point less than 0.2 of the lysosomes fraction colocalized with NP, which is a lower amount compared to the 4 h time-point **(Figure 4C)**.

**Figure 4.**
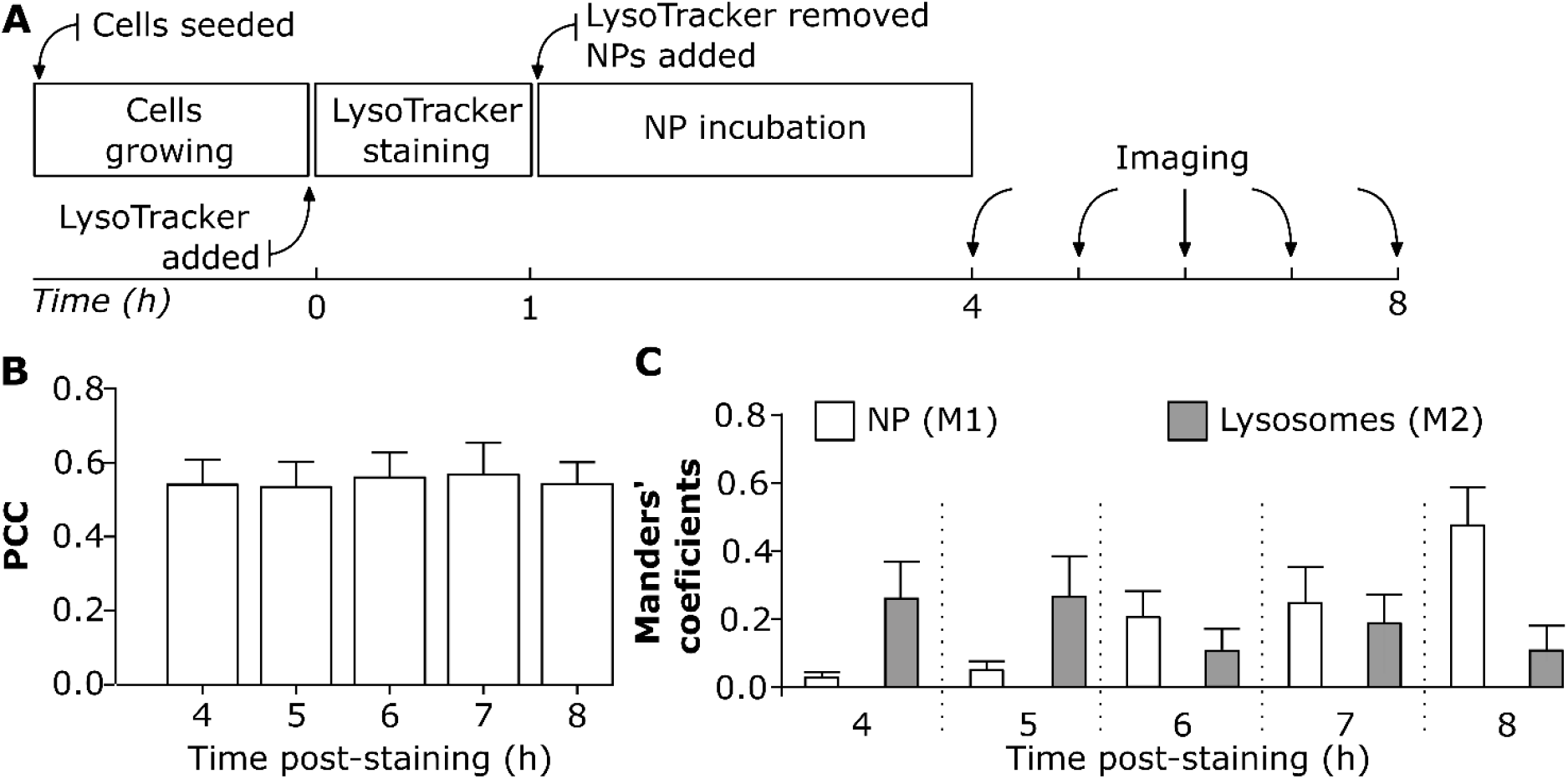
The effect of continuous live cell imaging on NP colocalization with lysosomes. **A**. Experimental workflow: Cells were first stained with LysoTracker Red probe for 1 h, and then exposed to 20 μg/mL of 59 nm SiO_2_-BDP FL NP in phenol free cRPMI. Cells were imaged continuously from 4 h to 8 h. All the images were analyzed using ImageJ with a script (script SI2). Each cell was analyzed individually. **B**. Pearson correlation coefficient (PCC) demonstrating correlation between NP and LysoTracker Red probe (lysosomes) at different time points. **C**. Manders’ correlation coefficient M1 corresponds to the fraction of 59 nm SiO_2_-BDP FL NP within lysosomes (stained with LysoTracker Red) and M2 corresponds to the fraction of lysosomes filled with NP. The corresponding histograms of fluorescence intensities of each channel is shown in **Figure S5A**.

Based on different NP physicochemical properties such as core material, size, shape, surface charge, porosity, surface roughness, surface area, crystalline structure, NP uptake and their intracellular fate can vary over time. In order to investigate these phenomena, we monitored the changes in colocalization of NP of different sizes and incorporated fluorophores.

#### Experiment II: The effect of different particles sizes on colocalization with lysosomes

A combination of the PCC and Manders’ colocalization coefficients can provide different messages of NP accumulation in lysosomes.^40^ Since particles of different sizes are widely used in the research, we chose 119 nm and 920 nm fluorescently labelled SiO_2_ particles labelled with the Cyanine5 (Cy5) fluorophore and tested their colocalization with lysosomes. Particles showed similar zeta potentials (−44 mV). After J774A.1 exposure to a single particle type for 24 h, cells were washed, stained with LysoTracker Red probe for 1 h and then visualized by live cell imaging **(Figure 5A)**.

**Figure 5.**
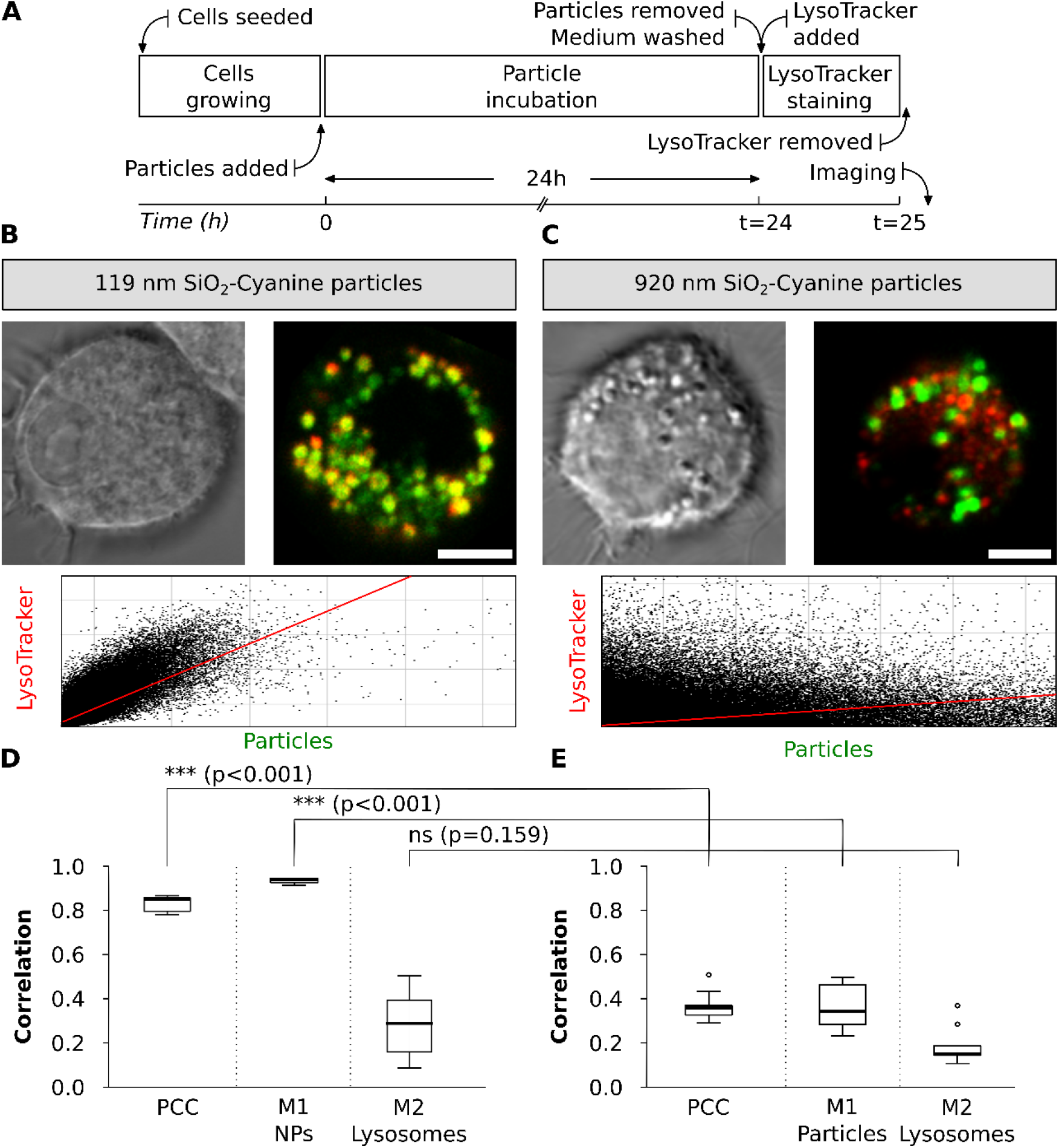
Colocalization of different-sized SiO_2_ particles with lysosomes in J774A.1 macrophages. **A**. Experimental workflow: Cells were exposed to SiO_2_ particles of different sizes for 24 h followed by LysoTracker Red staining for 1 h and imaged live. **B**. Representative bright field and confocal images of single cells with the corresponding cytofluorograms show colocalization of 119 nm SiO_2_-Cy5 particles (green) with lysosomes (red). **C**. Colocalization of 920 nm SiO_2_-Cy5 particles (green) with lysosomes (red). Co-localized pixels appear in yellow. Scale bar: 10 μm. The corresponding histograms of fluorescence intensities of each channel is shown in **Figure S5A** and the bright field images identifying cell morphology in **Figure S6**.Quantification of colocalization of **D**. 119 nm SiO_2_-Cy5 particles and **E**. 920 nm SiO_2_-Cy5 particles with LysoTracker Red probe was performed using the ImageJ software with JACoP plugin (n= 10 cells). Data is presented as Pearson’s correlation coefficient (PCC) and Manders’ overlap coefficients, where M1 represents the fraction of particles-associated pixels overlapping with lysosomes and M2 the fraction of lysosome-associated pixels overlapping with particles. Statistical analysis was performed using unpaired t-test in GraphPad Prism software: *** p < 0.001, ns – not significant.

High accumulation of 119 nm SiO_2_-Cy5 particles within lysosomes was observed after 24 h of particle exposure as demonstrated by the PCC of 0.9. This suggests the particles uptake and intracellular sorting to the lysosomes. Such information cannot be extracted by the naked eye from the CLSM images but can be visualized by the cytofluorograms **(Figure 5B):** the point cloud expanded towards the top right corner, and the increase in PCC reflected this process. As represented by Manders’ coefficients, almost all internalized 119 nm SiO_2_-Cy5 particles were associated with the lysosomes (M1 > 0.9) after 24 h. In addition, there were still lysosomes available without particles, as demonstrated by 0.3 of lysosome fraction (M2, **Figure 5D**).

A different outcome was observed with the 920 nm SiO_2_-Cy5 particles **(Figure 5 C, E)**. Only 0.3 of the particles fraction arrived to the lysosomes (PCC: 0.3) at 24 h upon particle exposure. Manders’ coefficients show that only 0.4 of particles fraction reached lysosomes within 24 h (M1: 0.4) and the majority of lysosomes fraction (M2: 0.2) were still left without particles (M2, **Figure 5E**). Thus, micron-sized particles were less likely to enter the lysosomes within 24 h and possibly sedimented in the cells’ surroundings or accumulated in other intracellular compartments.^65–67^ A possible explanation for lower colocalization of bigger particles with lysosomes could be as well the differences in particle number added to the cells (9×10^9^ of 119 nm SiO_2_-Cy5 particles per mL vs 2×10^7^ of 920 nm SiO_2_-Cy5 particles per mL) or, alternatively, different mobility of particles within the cells.^68^

#### Experiment III: Colocalization between NP with different fluorophores and lysosomes

A potential colocalization pitfall is the dependence of fluorescent dye encapsulation on the physicochemical properties of NP. The dye encapsulation might change the physicochemical properties of NP and the fluorescence properties of a dye, for example, by inducing fluorescence quenching or enhancing emission.^26,69,70^ An experimental design was implemented to compare the colocalization of NP with similar sizes but different fluorophores. J774.A1 cells were initially exposed to 59 nm SiO_2_-BDP FL NP and 59 nm SiO_2_-Rhodamine B (RhoB) NP for a period of 13 h. The medium containing NP was subsequently removed and cells were washed. Next, a fresh medium without NP was added and a resting phase of 6 hours was performed to investigate whether all the NP that have just reached the cell membrane continued being internalized and accumulate in the lysosomes. The resting time was chosen based on known literature, as it has been shown that different NP or macromolecules reach the lysosomes in a period from 3 h to 6 h.^20,71–75^ Finally, the cells were fixed and stained with LAMP-2 conjugated with a secondary antibody Alexa Fluor 647, which was selected due to the possible combination with NP-conjugated fluorophores **(Figure 6A)**. The cells exposed to 59 nm SiO_2_-BDP FL NP and 59 nm SiO_2_-RhoB NP were stained with LAMP-2 and imaged **(Figures 6B and 6C)**.

**Figure 6.**
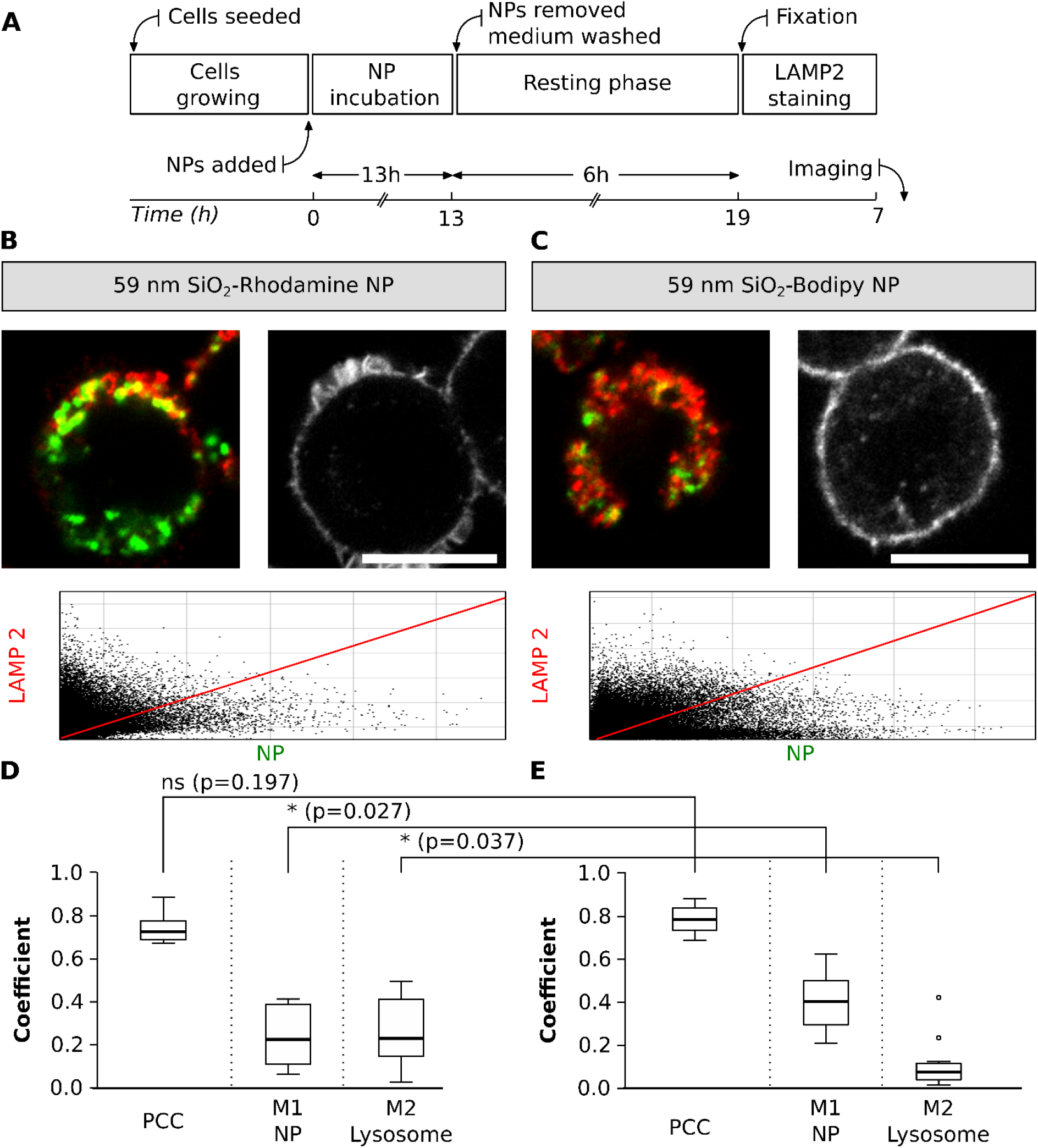
Colocalization between NP with different fluorophores and lysosomes. **A**. Experimental workflow. Representative images of single cells exposed to two different SiO_2_ NP (green), **B**. SiO_2_-Rho NP and **C**. SiO_2_-BDP FL NP. Lysosomes were stained with LAMP-2 antibody (shown in red). Scale bar: 10 μm. The corresponding histograms of fluorescence intensities of each channel is shown in **Figure S5B** and the F-actin staining identifying cell morphology in **Figure S6**. The plots represent main parameters (PCC, M1 and M2) to estimate colocalization between different NP and lysosomes. The colocalization between the **D**. 59 nm SiO_2_-RhoB NP and **E**. 59 nm SiO_2_-BDP FL NP with lysosomes, labelled with LAMP-2 antibody. PCC: Pearson correlation coefficient, Manders’ correlation coefficients M1 and M2. The whiskers represent the standard deviations. Analysis was performed in individual cells for each experiment (n=10-13 cells). All the images were analysed using ImageJ, with the JACoP plugin comparing the correlation coefficients. Statistical analysis was performed using unpaired t-test in GraphPad Prism software. * p < 0.05, ns – not significant.

Based on PCC values, the results showed no significant differences in the colocalization of lysosomes and NP with different fluorophores (PCC: 0.8). Nevertheless, Manders’ correlation coefficients (M1, fraction of NP inside the lysosomes) confirmed that 0.2 of SiO_2_-RhoB NP fraction are localized in the lysosomes **(Figure 6D)** compared to 0.4 of SiO_2_-BDP FL NP fraction **(Figure 6E)**. Additionally, a different fraction of lysosomes filled with NP (M2) was observed when comparing NP with different fluorophores, 0.3 for the cells exposed to SiO_2_-RhoB NP and 0.1 for the cells exposed to SiO_2_-BDP FL NP. This could be explained by the difference in zeta potential between SiO_2_-BDP FL NP (−54 mV) and SiO_2_-RhoB NP (−34 mV). As reported previously in literature, NP uptake and their intracellular fate are strongly influenced by the NP surface properties.^18,70,76–78^

#### Experiment IV: Comparison of NP colocalization with lysosomes between fixed-and live-cell imaging

While performing lysosomal staining, both LysoTracker probes and LAMP antibodies are available. However, there are differences both in their specificity as well as in staining protocols, which can significantly impact the experimental outcome.^28^ With this experimental approach, the changes in colocalization of NP with lysosomes between live- and fixed-cell imaging were evaluated. The same exposure time and resting phase was kept as described in section III **(Figure 7A)**. In addition to the LysoTracker staining, cells were immunostained with LAMP-2 antibody **(Figure 7A, 7B and 7D)** and the results from fixed-cells were compared to the data collected in section III.

**Figure 7.**
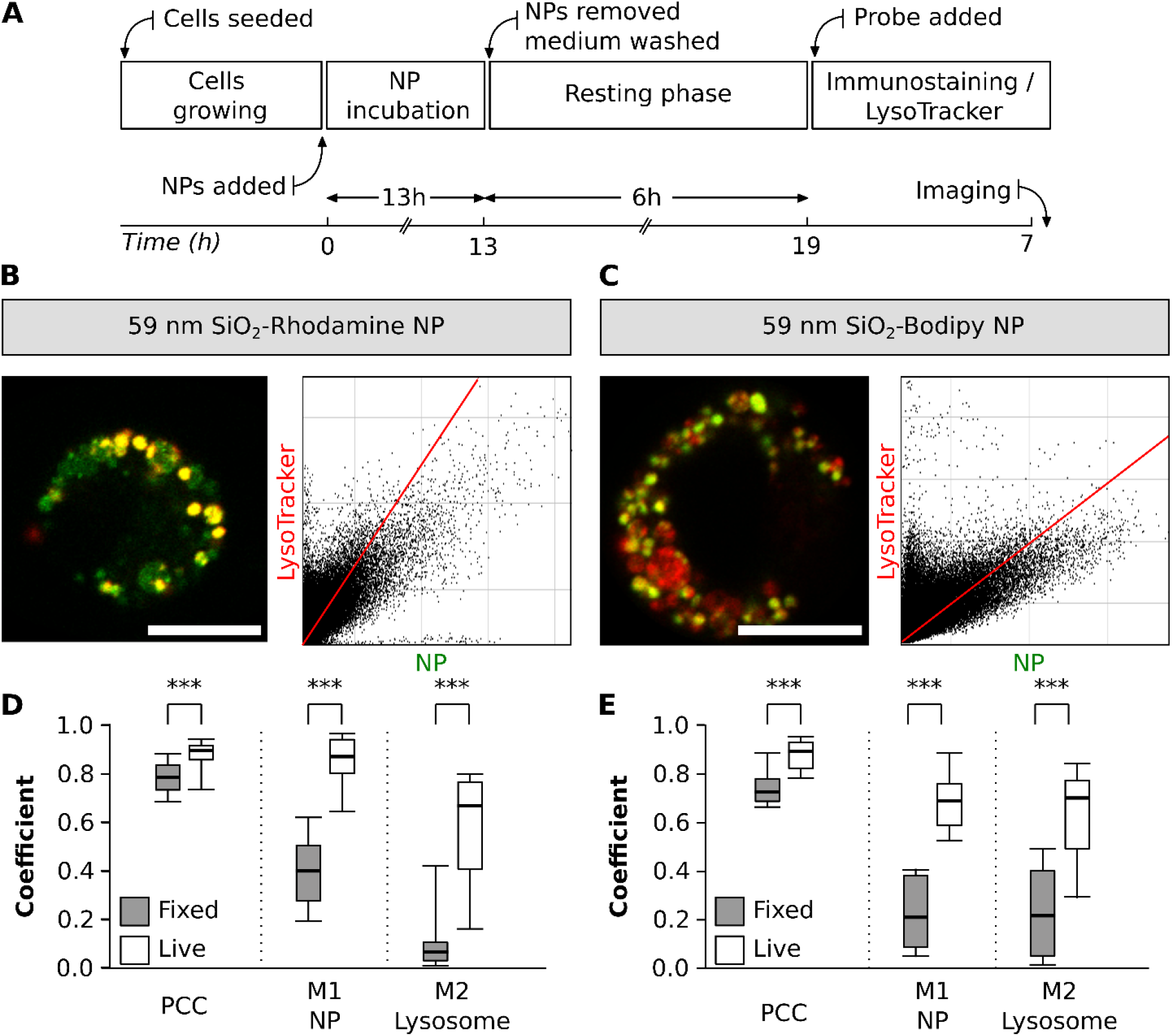
NP colocalization with lysosomes between fixed-and live-cell imaging. **A**. Experimental workflow. Representative images of a single cell exposed to two different NP types, **B**. SiO_2_-RhoB NP (green) and **C**. SiO_2_-BDP FL NP (green). Lysosomes were stained with LAMP-2 antibody (red). The corresponding histograms of fluorescence intensities of each channel is shown in **Figure S5B**. The graphs depict the main parameters (Pearson correlation coefficient - PCC, Manders’ correlation coefficients - M1 and M2) to estimate the colocalization between **D**. SiO_2_-RhoB NP and lysosomes (LysoTracker Green) and **E**. SiO_2_-BDP FL NP and lysosomes (LysoTracker Red) in live cells (white box) and fixed cells stained with and LAMP-2 (grey box). The whiskers are the standard deviations. The analysis was done in individual cells for each experiment (n=8-13 cells). All the images were analysed using ImageJ with the JACoP plugin comparing the correlation coefficients. Statistical analysis was performed using unpaired t-test in GraphPad Prism software. *** p < 0.001, ns – not significant.

No significant changes in PCC values were observed among live and fixed cells for both NP types. Nevertheless, we observe differences for Manders’ correlation coefficients (M1 and M2) **(Figure 7C and 7E)**. In live cells, exposed to 59 nm SiO_2_-BDP FL NP, there were roughly 0.7 of NP fraction colocalized with the lysosomes (M1), but in fixed-cell the colocalization decreased to 0.4 **(Figure 7C)**. This difference is evident in the cells exposed to 59 nm SiO_2_-RhoB NP where 0.9 of the NP are colocalized with the lysosomes of living cells stained with LysoTracker in contrast to only 0.2 of colocalized NP inside the lysosomes of fixed-cells **(Figure 7E)**. Likewise, a decrease in the fraction of lysosomes filled with NP (M2) between live and fixed cells was noted. These differences are attributed to image processing, and selectivity of LysoTracker probe (stains lysosomes and late endosomes) vs LAMP-2 (specific to LAMP-2 protein present on lysosomal membrane).^79^

Correlative studies between live and fixed cells are challenging because the shape of the fixed cells varies from its natural state.^80^ Moreover, a quantitative analysis that compares and correlates the intensities of fluorophores such as the LysoTracker probes after cell fixation is expected to be less precise due to the decrease of LysoTracker intensity and further changes in cellular shape **(Figure S7 and Figure S8)**. We have already described that each experimental factor that alters the intensity of the fluorophores will have repercussions, especially on the Manders’ coefficients, as shown in **Figure S8** and **Figure S9**. In conclusion, many parameters influence the NP-lysosome colocalization outcome such as the fluorophore dye to label the NP, the staining procedure for lysosomes in cells, the lysosome marker specificity, and the imaging set-up. Therefore, it is recommended to consider these parameters when planning and conducting the experiments as well as to adjust each experiment individually based on the research question to minimize the variability.

## 3. Conclusions

We visualized the lysosomes by labeling cells with LysoTracker probes and calculated the colocalization between fluorescently labelled NP and lysosomes using different correlation coefficients. Herein, we described detailed steps and recommendations from the experimental design, staining, sample preparation and imaging on the colocalization parameters.

Key considerations for conducting NP-lysosome imaging and colocalization analysis are:

i. Fluorescent lysosomal probe properties (see Table 1), such as concentration to be used, incubation time and fluorophore,^18^
ii. Physicochemical characteristics of the NP (see Table 2) such as exposure time, concentration, and administration method,^81^
iii. CLSM imaging setup including resolution, illumination, and storage considerations,^82,83^
iv. Colocalization analysis of NP in the lysosomes.^36^

We strongly advise to fully report the parameters of the image acquisition to allow correct comparisons and reproducibility between experiments reported from different studies. The human eye is not capable of extracting the correct colocalization interpretation from CLSM images alone: a cytofluorogram must be added. Similarly, a single PCC often does not provide information about the fractional distribution of the fluorophores co-occurring within the resolution of a single voxel, for this an additional measure such as Manders’ co-occurrence coefficients should be taken into analysis. Additional recommendations are summarized in the **Table 3**.

**Table 3.**
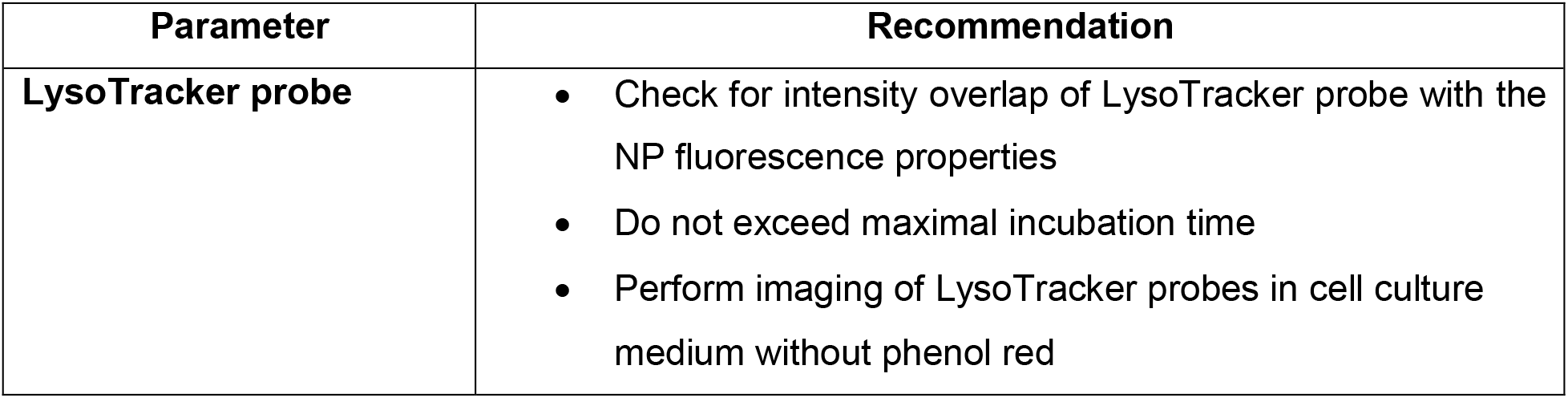
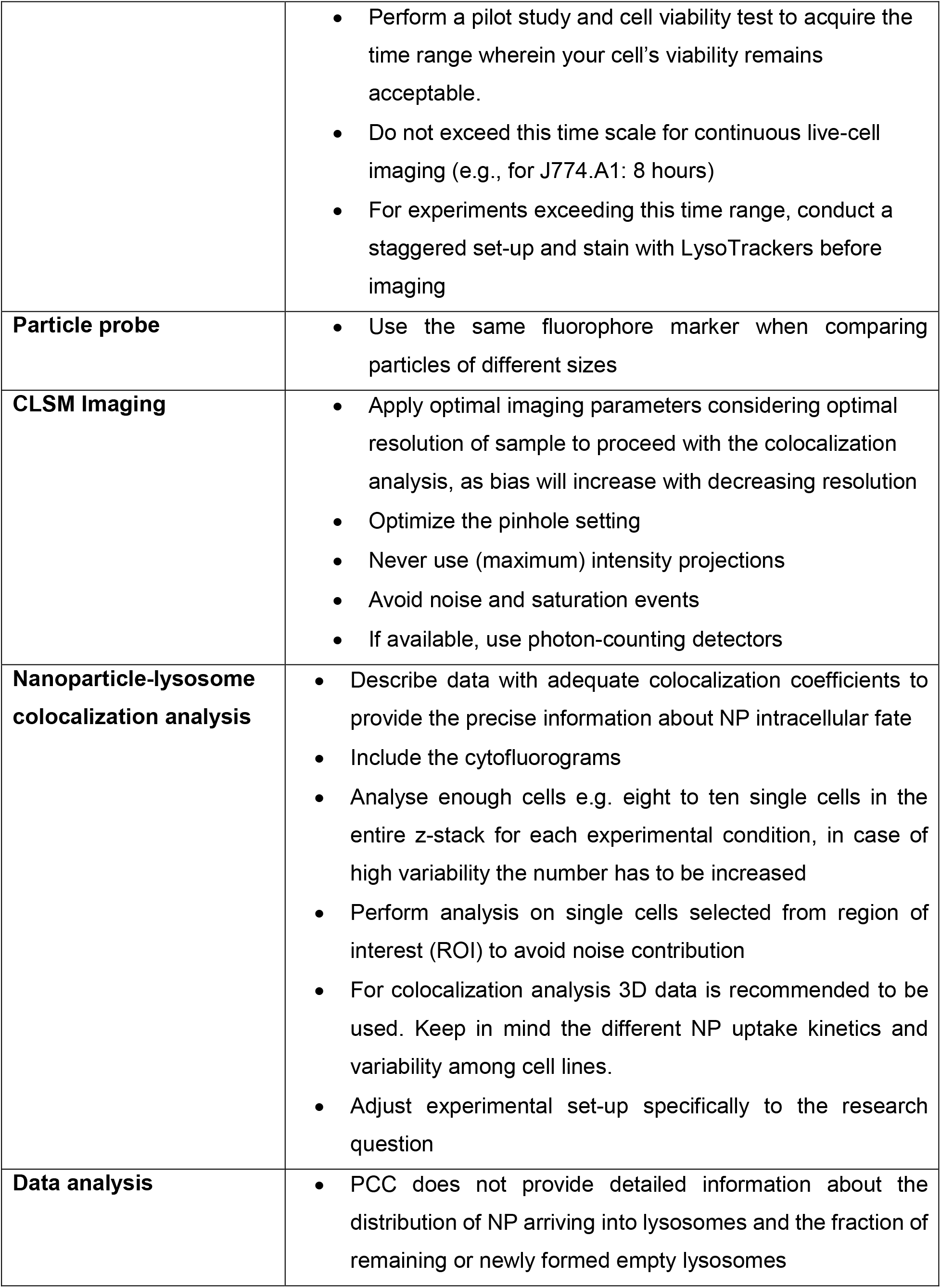
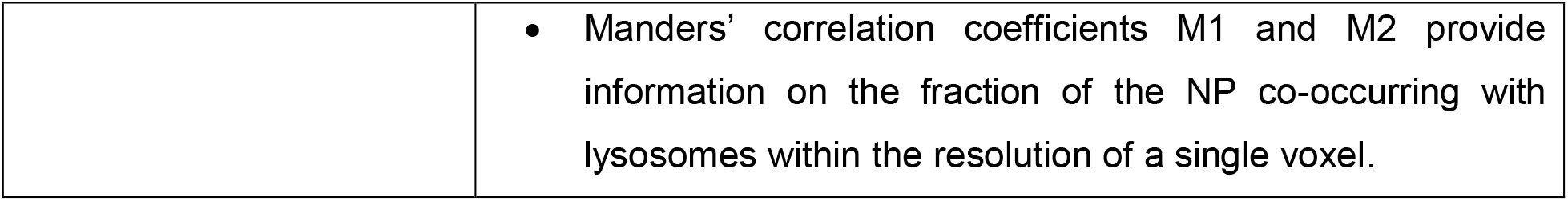
List of recommendations based on the different parameters.

| Parameter | Recommendation |
| --- | --- |
| LysoTracker probe | • Check for intensity overlap of LysoTracker probe with the NP fluorescence properties<br>• Do not exceed maximal incubation time<br>• Perform imaging of LysoTracker probes in cell culture medium without phenol red |
|  | • Perform a pilot study and cell viability test to acquire the time range wherein your cell's viability remains acceptable.<br>• Do not exceed this time scale for continuous live-cell imaging (e.g., for J774.A1: 8 hours)<br>• For experiments exceeding this time range, conduct a staggered set-up and stain with LysoTrackers before imaging |
| **Particle probe** | • Use the same fluorophore marker when comparing particles of different sizes |
| **CLSM Imaging** | • Apply optimal imaging parameters considering optimal resolution of sample to proceed with the colocalization analysis, as bias will increase with decreasing resolution<br>• Optimize the pinhole setting<br>• Never use (maximum) intensity projections<br>• Avoid noise and saturation events<br>• If available, use photon-counting detectors |
| **Nanoparticle-lysosome colocalization analysis** | • Describe data with adequate colocalization coefficients to provide the precise information about NP intracellular fate<br>• Include the cytofluorograms<br>• Analyse enough cells e.g. eight to ten single cells in the entire z-stack for each experimental condition, in case of high variability the number has to be increased<br>• Perform analysis on single cells selected from region of interest (ROI) to avoid noise contribution<br>• For colocalization analysis 3D data is recommended to be used. Keep in mind the different NP uptake kinetics and variability among cell lines.<br>• Adjust experimental set-up specifically to the research question |
| **Data analysis** | • PCC does not provide detailed information about the distribution of NP arriving into lysosomes and the fraction of remaining or newly formed empty lysosomes |
|  | • Manders' correlation coefficients M1 and M2 provide information on the fraction of the NP co-occurring with lysosomes within the resolution of a single voxel. |

## 4. Methods

### 4.1. Synthesis of silica nanoparticles

The synthesis of fluorescently labelled silica (SiO_2_) NP was performed following a Stöber method.^84^ For a detailed description of the 59 nm SiO_2_-BDP FL NP and 920 nm SiO_2_-Cy5 NP synthesis and characterization, refer to the publication from Susnik et al.^32^ Similar synthesis protocol was adapted for 59 nm SiO_2_-RhoB NP using a RhoB isothiocyanate (RBITC) dye (CAS 36877-69-7, Sigma-Aldrich, USA). For 119 nm SiO_2_-Cy5 NP synthesis, briefly 58 mL of water (Milli-Q), 162 mL of absolute ethanol (EtOH) (VWR, Dietikon, Switzerland), and 8.71 mL of 25% ammonium hydroxide (NH4OH) (Merck, Zug, Switzerland) were heated at 70 °C in a 500 mL flask and sealed. After 30 min, 22 mL of tetraethyl orthosilicate (TEOS) were added to the mixture. Finally, after 2 min, a mixture of 200 μL-Cy5 NHS ester (11.5 mg/mL in dimethyl sulfoxide (DMSO), Lumiprobe 53020, Hunt Valley, MD, USA) and 3 μL of (3-aminopropyl) triethoxysilane (APTES) were added to the flask. The reaction was further heated for 4 h, cooled down to room temperature, and centrifuge at 2000 rpm for 40 minutes three times and redisperse in Milli-Q water. The obtained particles in dispersion were stored in the dark at 4 °C. **Table 2** summarizes the physicochemical characteristics of three different SiO_2_ particles, such as core diameter measured by transmission electron microscopy (TEM; FEI Technai G2Spirit, Thermo Fisher Scientific, Waltham, MA, USA). Hydrodynamic diameter of NP in Milli-Q water and cRPMI was measured by dynamic light scattering (DLS) at constant temperature (37 °C) using a commercial goniometer instrument (LS Spectrometer, LS Instruments AG, Switzerland) equipped with a linearly polarized and a collimated laser beam (Cobolt 05-01 diode-pumped solid-state laser, λ = 660 nm, P max. = 500 mW). Data were collected at scattering angle of 90° and laser wavelength 660 nm. Two APD detectors, assembled for pseudo-cross-correlation, were used to improve the signal-to-noise ratio. The mean and standard deviation were calculated from ten independent measurements. The surface charge was determined by phase-amplitude light scattering (ZetaPALS, Brookhaven Instruments Corp., Holtsville, NY, USA) in Milli-Q water.

### 4.2. Cell culture and seeding

Mouse monocyte/macrophages J774A.1, obtained from the American Type Culture Collection (ATCC, Rockville, MD, USA) were subcultured Roswell Park Memorial Institute 1640 medium (RPMI; Cat.No. 11835030, Gibco, Life Technologies, Zug, Switzerland) supplemented with 10 % v/v FBS, 1 % v/v penicillin/streptomycin and 1 % v/v L-glutamine. Cells were maintained in T75 polystyrene cell culture flasks (TRP, Trasadingen, Switzerland) in cRPMI until reaching 80 % confluence. Cell seeding procedure has been performed as described by Susnik et al.^32^

### 4.2. The interference of phenol red with the LysoTracker probes

LysoTracker™ Red DND-99 (Cat. No. L7528), Green DND-26 (Cat. No. L7526) and Blue DND-22 (Cat. No. L7525) (Thermo Fisher Scientific, MA, USA) probes at final concentration 75 nM were prepared in cRPMI medium containing phenol red or in phenol red-free medium. J774A.1 macrophages were incubated with LysoTracker probes for 1 h at 37 °C, 5 % pCO_2_ and 95 % relative humidity, following providers’ protocol (Invitrogen, Thermo Fisher Scientific, Waltham, MA, USA). After incubation, the medium containing LysoTracker probes was removed and replaced with fresh phenol red-containing or phenol red-free medium. Live cell imaging was done in cRPMI containing phenol red or in phenol free medium (refer to chapter Image acquisition). The images were analyzed using ImageJ software with a JACoP plugin and a custom-written script **(Figure S2** and **Script S1)**. Briefly, each cell contour was manually defined based on the LysoTracker channel to define the region of interest (ROI) for the analysis. The detected signals from the LysoTracker channel were masked, followed by an automatically adjusted threshold. An inverted mask was used to remove the background noise and to measure the raw integrated density by excluding background signals. The raw integrated densities for ten individual cells were analyzed for each LysoTracker probe. The following parameters have been used: 16-bit image (1024×1024 pixels), pixel dwell of 25.5,1 Airy unit and the gain for each individual fluorophore as it follows: LT blue 850, pinhole 43, channel range 410-431 nm; LT green 583 pinhole 52, channel range 493-606 nm; LT Red 446, pinhole 60, channel range 566-690 nm.

### 4.3. Stability of LysoTracker Red probe

For continuous imaging set-up, LysoTracker™ Red DND-99 (Cat. No. L7528), (Thermo Fisher Scientific, MA, USA) probe at final concentration 75 nM was prepared in cRPMI medium containing phenol red or in phenol red-free medium. J774A.1 macrophages were incubated with LysoTracker probes for 1 h at 37 °C, 5 % pCO_2_ and 95 % relative humidity, following providers’ protocol (Invitrogen, Thermo Fisher Scientific, Waltham, MA, USA). After incubation, the medium containing LysoTracker probes was replaced with fresh phenol red-free medium without LysoTracker probe. Cells were immediately imaged live (refer to chapter Image acquisition) in a continuous (24h) or staggered set-up (1h, 4h, 24h). The images were analyzed using ImageJ software with a JACoP plugin and a custom-written script as in described previously. The raw integrated densities for individual cells were analyzed. The following parameters have been used: 16-bit image (1024×1024 pixels) and pixel dwell of 25.5,1 Airy unit.

### 4.4. Cell viability

Cells were incubated with LysoTracker Red probe for 1 h, 4 h and 24 h under standard cell culture conditions. Supernatants were collected and cell viability was assessed based on the lactate dehydrogenase (LDH) enzyme released from the damaged cells. Data was analyzed in triplicate using LDH cytotoxicity detection kit (Roche Applied Science, Mannheim, Germany) according to the manufacturer’s protocol. The absorbance of colorimetric product formazan was recorded using a microplate reader (Bio-Rad, Switzerland) at 490 nm with a reference wavelength of 630 nm. The values were expressed as fold increase of the treated samples relative to the untreated (control) samples. Triton X-100 (0.2% v/v; in phosphate buffered saline (PBS)) (Merck, Switzerland) was added to the positive control 15 min prior to collecting the supernatant of each well.

### 4.5. Cell exposure to silica NP and LysoTracker probes

A suspension of 1 mg/mL silica NP was first prepared in Milli-Q water. NP were dispersed in cRPMI to the final concentration of 20 μg/mL and administered to the cells via a pre-mixed method i.e., NP were added to cRPMI immediately prior to the cell exposure. Three experimental setups were designed based on the addition of the LysoTracker probes:

i. **Addition of LysoTracker probe before NP exposure for live cell imaging:** Cells were seeded in uncoated glass-bottom MatTek culture dishes (35 mm, Life Sciences, Massachusetts, USA) for 24 h and incubated with fresh cRPMI supplemented with 75 nM LysoTracker Red probe for 1 h at 37 °C. Afterwards, the supernatant containing LysoTracker Red probe was removed, and cells were exposed to 59 nm SiO_2_-BDP FL NP in cRPMI for a period of 8 h. The image acquisition was performed from 4 h to 8 h in a chamber at 37 °C and 5 % pCO_2_, adapted for the Zeiss LSM710 confocal microscope. The cells were imaged as hyperstacks over time. Time lapse and different z-stacks were set for each sample. The time point was defined upon NP exposure and the analysis was done from 4 h to 8 h. For each time point, 13 individual cells were analyzed to evaluate the raw integrated intensity of LysoTracker probe over time. The raw integrated density was processed following the same procedure as described in section 4.3. The correlation coefficients were calculated using ImageJ software with JACoP plugin and a custom-written script (see **Script S2**).
ii. **Addition of the LysoTracker probe after the NP exposure:** Cells were initially exposed to a single particle type for 24 h. The cells were washed 3-times with 1X PBS and incubated with fresh cRPMI supplemented with 75 nM LysoTracker probe for 1 h (according to the manufacturers’ recommendations). Afterwards, the supernatant containing LysoTracker probe was removed, and fresh cRPMI was added to the cells for immediate live-cell imaging.
iii. **Addition of the LysoTracker probe after NP incubation and followed by resting period:** Cells were exposed to 59 nm SiO_2_-RhoB NP or 59 nm SiO_2_-BDP FL NP for 13 hours, washed 3-times with 1X PBS and further incubated with fresh cRPMI. Cells were incubated for an additional 6 h to ensure that most of the NP reached the lysosomes. After, cells were stained with 75 nM LysoTracker Green or 75 nM LysoTracker Red for 60 minutes. The LysoTracker probes were replaced with fresh cRPMI and cells underwent immediate live-cell imaging.

### 4.6. Image acquisition

During live-cell imaging, cells were kept in phenol-red containing cRPMI or alternatively in cRPMI without phenol red. Image acquisition was performed with Zeiss LSM 710 Meta confocal microscope Axio Observer.Z1 (Zeiss GmbH, Jena, Germany), using a Plan-Apochromat objective 63×/NA 1.4-immersion oil objective lens fitted with diode-pumped solid-state laser (DPSS) 561-10, argon 458, 488, 514 and HeNe 633 nm lasers. An immersion oil (Carl Zeiss Immersol™ 518F) with a refractive index 1.518 was used. The excitation/emission spectra peaks for LysoTracker probes: Blue (λ_ex_: 373 nm; λ_em_: 422 nm), Green (λ_ex_: 504 nm; λ_em_: 511 nm) and Red (λ_ex_: 577 nm; λ_em_: 590 nm), using the respective lasers (405 nm, argon 458 nm and DPSS 561 nm) with adjusted master gain and pinhole for each of the experiments. The excitation/emission spectra peaks for NP are the following: SiO_2_-BDP FL NP (laser λ_ex_: 488 nm; λ_em_: 520/30 nm), SiO_2_-RhoB NP (λ_ex_: 561 nm; λ_em_: 620/30 nm), and SiO_2_-Cy5 NP (λ_ex_: 633 nm; λ_em_: 700/60 nm). After acquisition, the images were analyzed using FIJI software (ImageJ, National Institutes of Health, Bethesda, MD, USA).

### 4.7. Colocalization analysis

To quantify the degree of colocalization between fluorescently labeled silica NP and LysoTracker probes, the Pearson correlation coefficient (PCC) and the Manders’ correlation coefficient (M1 and M2) were determined from the corresponding confocal images using FIJI software (ImageJ, National Institutes of Health, Bethesda, MD, USA) with JACoP plugin and a custom-written script **(Script S2)**. We used an approach without Costes randomizations, taking the automatic adjusted threshold for the two different channels. The cell contours were manually defined based on the LysoTracker channel to select the region of interest (ROI) for the analysis and avoid background signals. In the script run for each channel the automatic threshold was adjusted, where channel 1 represented nanoparticles and channel 2 corresponded to a LysoTracker probe. All the images were analyzed with the same script and the threshold of each individual channel was adjusted each time. From the JaCoP analysis we extracted Pearson and Manders’ (M1 and M2) correlation coefficients for data representation.

### 4.8. Data post-processing and simulation

J774A.1 macrophages were exposed to 59 nm SiO_2_-Bodipy fluorescein NHS ester (SiO_2_-BDP FL) NP and 59 nm SiO_2_-RhoB NP for 13 h. Cells were washed to remove the remaining NP, followed by resting phase fixation and immunostaining for LAMP-2 and subsequently recorded on a Zeiss LSM 710 meta laser scanning laser microscope.

Post-recording, two channels of the dataset were treated to mimic different imaging setups. The simulations were conducted in Fiji software and colocalization was performed in JaCoP plugin. **i)** To mimic loss of spatial resolution, the dataset was binned in XY using the build-in scaling function of Fiji. **ii)** Loss of axial resolution (increase in optical slice thickness) was created by the sum of slices of the grouped Z-project routine. **iii)** Pixel saturation was simulated by iteratively multiplying the intensity of each pixel by a factor 1.1 of an oversaturation-free stack and calculating the colocalization measures after each step. **iv)** The loss of bit depth was made by converting the original 16-bit stack to 8-bit and repeating the colocalization analysis. This part is divided in two segments: **i)** Immunostaining: Upon completion of the NP exposures, cells were washed with PBS and fixed with 4 % paraformaldehyde (in PBS, v/v) for 15 minutes. After washing, cells were blocked with 4 % bovine serum albumin (BSA) (in PBS, v/v) for 1 hour, washed with PBS and stained overnight at 4 °C with 20 μg/mL of LAMP-2 primary antibody (Thermo Fisher Scientific, MA, USA) in 1 % BSA (in PBS, v/v). After washing, Alexa 647-conjugated secondary antibody was applied (1:500 dilution) for 1 hour. Cells were washed and mounted (Fluoromount™ Aqueous Mounting Medium (F4680-25mL, Sigma-Aldrich) before proceeding with imaging with Zeiss LSM 710 confocal microscope. LAMP-2 corresponding excitations were for Alexa 647 varying the master gain and pinhole for each of the experiments. This data was used for the analysis of the colocalization coefficients for fixed-cells described in section VI-experiment III. **ii)** The simulations using scripts and macros (see SI): cLSM images were manipulated to evaluate the effect of spatial resolution, optical slice thickness, pixel saturation and bit depth on the colocalization parameters. The colocalization was analyzed in ImageJ software using the custom-written script and JACoP plugin. Representative data was selected to show the effect of different imaging conditions on the correlation coefficients.

### 4.10. Statistical Analysis

The statistical analyses were performed in GraphPad Prism 9 software, Origin 2016, 64 bit adjusting each of the treatments to a corresponding statistical analysis. One-way ANOVA, t-test or Dunnett’s test analysis were performed. The results are reported as the mean with standard deviation of the mean (SD).

## Abbreviations

NP: Nanoparticles
FIB-SEM: Focused ion beam-scanning electron microscopy
CLSM: Confocal laser scanning microscopy
LAMP: Lysosomal-Associated Membrane Protein
PCC: Pearson’s correlation coefficient
cRPMI: Complete Roswell Park Memorial Institute medium, supplemented with 10% fetal bovine serum, 1 % (v/v) L-glutamine and 1 % (v/v) penicillin/streptomycin
FBS: Fetal bovine serum
LDH: Lactate dehydrogenase
SiO_2_: Silica
DLS: Dynamic light scattering
M1 and M2: Manders’ coefficients
ROI: Region of interest
RBITC: Rhodamine B isothiocyanate
BSA: Bovine serum albumin
PBS: Phosphate buffered saline

## Supplementary information

**Figure S1A**. Comparison of the absorbance and emission spectra of the different LysoTracker probes. **Figure S1B**. Comparison of the emission spectra of the different (nano)particles. **Figure S1C**. Comparison of the absorbance spectra of the different (nano)particles. **Figure S2**. Confocal microscopy images of different LysoTracker probes. **Figure S3**. Transmission electron micrographs (TEM) of the different silica (nano)particles. **Figure S4A**. Size distributions of 59 nm SiO_2_-BDP FL NP and 59 nm SiO_2_-RhoB NP measured by dynamic light scattering. **Figure S4B**. Size distributions of 119 nm SiO_2_-Cy5 particles and 920 nm SiO_2_-Cy5 particles measured by dynamic light scattering. **Figure S5A and S5B**. Representative images of individual cells from different experiments for identification and contour with the corresponding histogram of the complete image acquired for each individual channel (nanoparticles and lysosomes). **Figure S6**. Representative bright field image of live cells **(A)** and fluorescent microscopy image of fixed cells **(B)** showing a population of individual cells used for colocalization analysis. **Figure S7**. Representative images of LysoTracker Red probe and Lamp-2 stainings in a single cell. **Figure S8**. Comparison of Pearson’s and Manders’ coefficients between live and fixed cells. **Figure S9**. Comparison of raw integrated intensity between live and fixed cells. **Video S1**. Continuous live cell imaging of J774A.1 cells. **Script S1**. Raw integrated densities. **Script S2**. Colocalization analysis.

## Acknowledgments

The authors would like to thank Liliane Ackermann Hirschi and Aaron Lee for supporting the synthesis of the nanoparticles.

## Author contributions

The influence of phenol red in cell culture medium on fluorescence intensity of LysoTracker probes and the stability of LysoTracker Red probe were tested by AMME. The analysis of nanoparticles within the lysosomes for the experiment I, III, IV was done by AMME and experiment II was done by ES. the simulations and post-processing imaging and the design of figure panels were done by DV. The support for the synthesis of the nanoparticles was done by PTB. The dynamic light scattering analysis was supported by SB. AMME and ES conducted the literature search, designed, and prepared a manuscript. AMME, ES, DV, BR, AK, DV participated in the writing, discussing and preparing the final manuscript. The supervision by BR and AK. The graphical abstract was done by AMME. All authors have read and agreed to the published version of the manuscript.

## Founding

The authors acknowledge funding from the Swiss National Science Foundation (SNSF) (project 310030_159847/1) and the Adolphe Merkle Foundation. Additionally, this work benefitted from support from the Swiss National Science Foundation through the National Center of Competence in Research Bio-Inspired Materials.

## Declarations

### Consent for publication

All authors have approved the submitted manuscript.

## Supplementary information

**Figure S1A.**
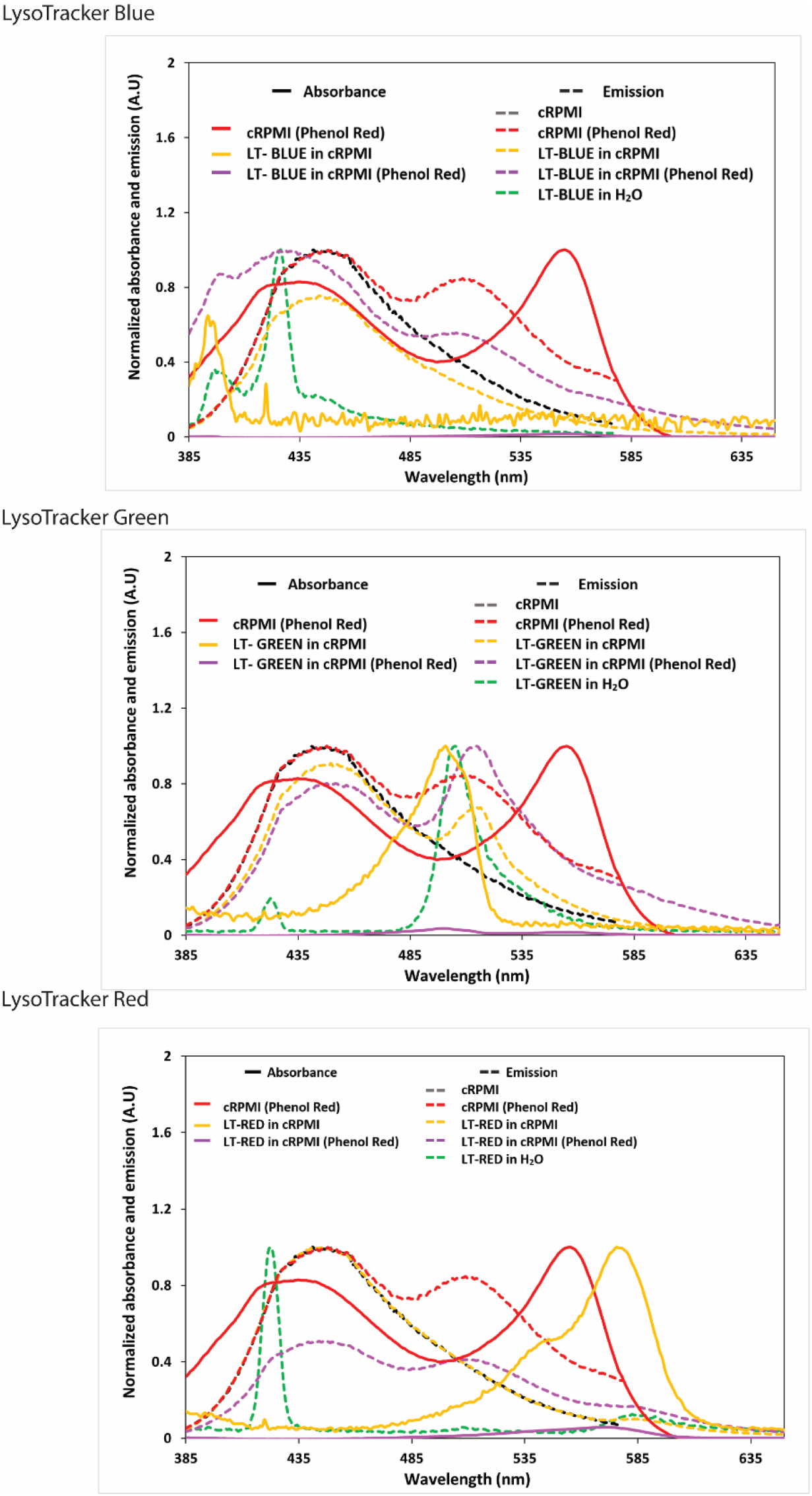
Comparison of the absorbance and emission spectra of the different LysoTracker (LT) probes, in cRPMI with or without phenol red and the control emission spectra in Milli-Q water. Lines: absorbance, dash lines: emission. Note: The UV-Vis Absorption Spectroscopy was carried out with a Jasco V-670 spectrophotometer. Quartz cuvettes (1 cm, science outlet) were used. The fluorescence Spectroscopy measurements were recorded with a Horiba Fluorolog 3 spectrometer equipped with a 450 W Xenon light source for excitation.

**Figure S1B.**
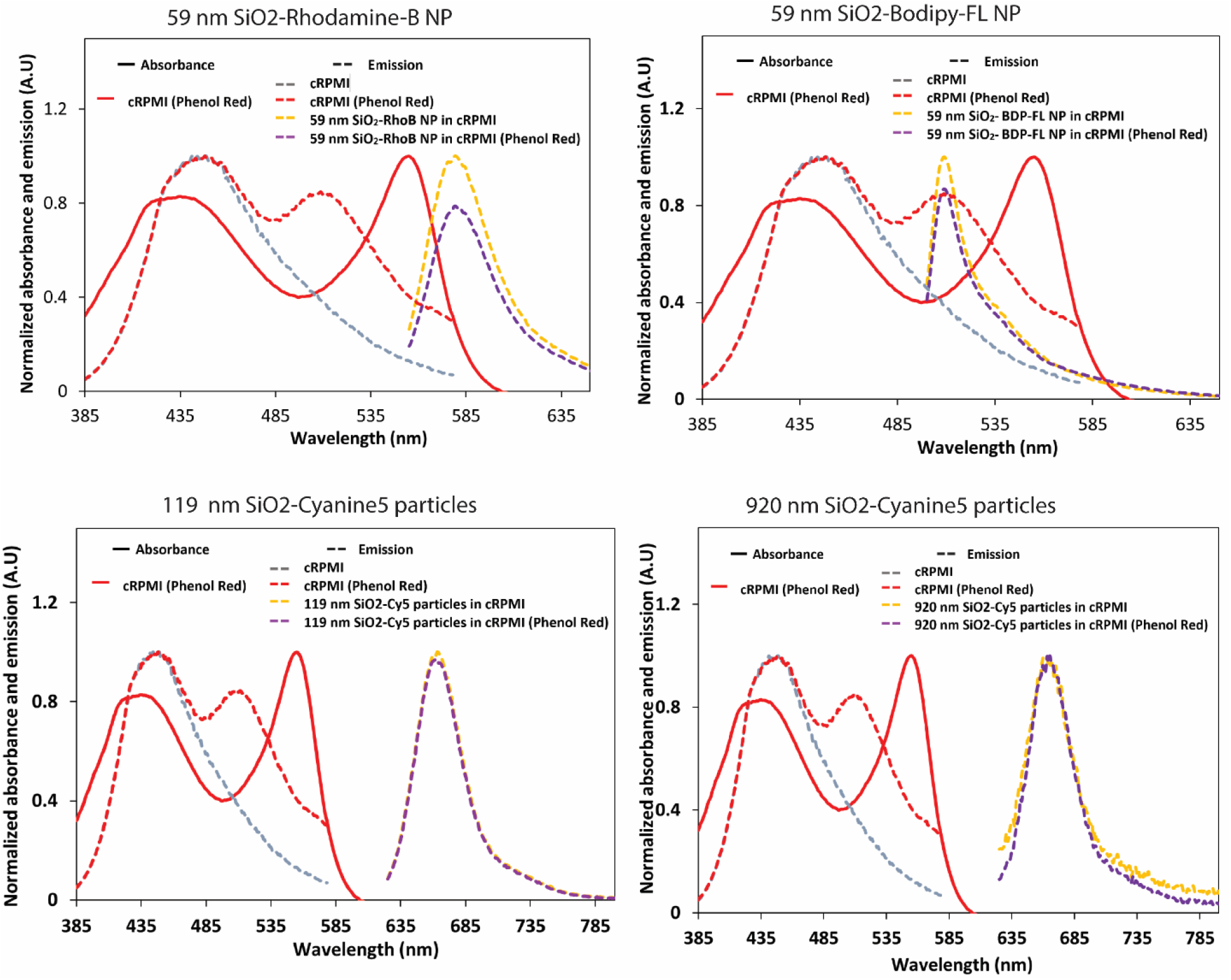
Comparison of the emission spectra of the different (nano)particles in cRPMI with and without phenol red and the absorbance of cRPMI with phenol red. Lines: absorbance, dash lines: emission. For the corresponding data to (nano)particles we measured the emission of (nano)particles instead of the pure dye. Since all the experiments were performed with the (nano)particles labeled with a specific dye, the microscopy fluorescent signals will correspond to the fluorophores that are incorporated into the (nano)particles. This makes the measurement closer to the real experimental conditions.

**Figure S1C.**
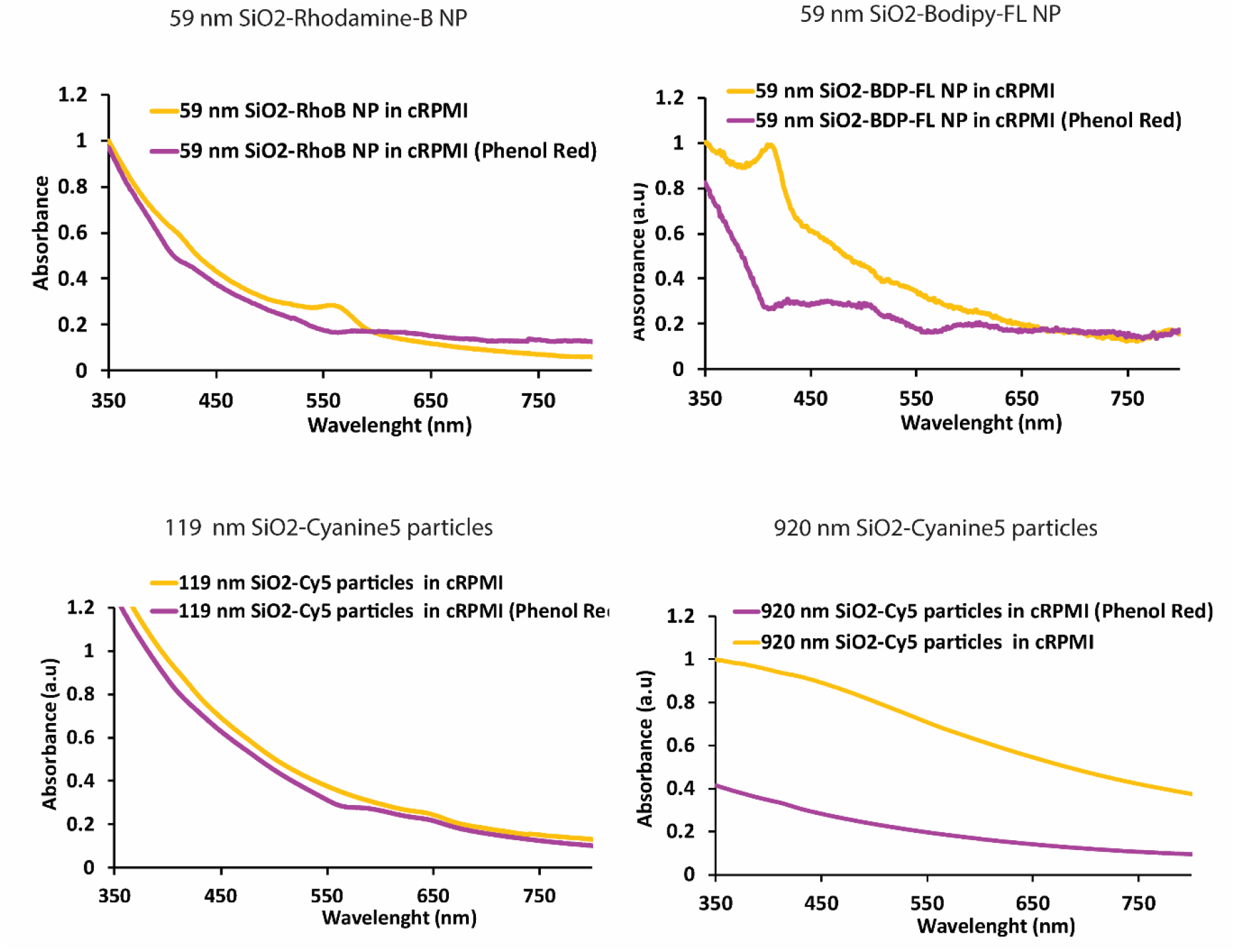
Comparison of the absorbance spectra of the different (nano)particles in cRPMI with and without phenol red. The absorbance spectra of the fluorophores in (nano)particles is dominated by the scattering of (nano)particles. This is the reason why the characteristic peak of the fluorophores is not strongly visualized. Nevertheless, the characteristic peak of the fluorophore is clearly visualized in the emission spectra **Figure S1 B**.

**Figure S2.**
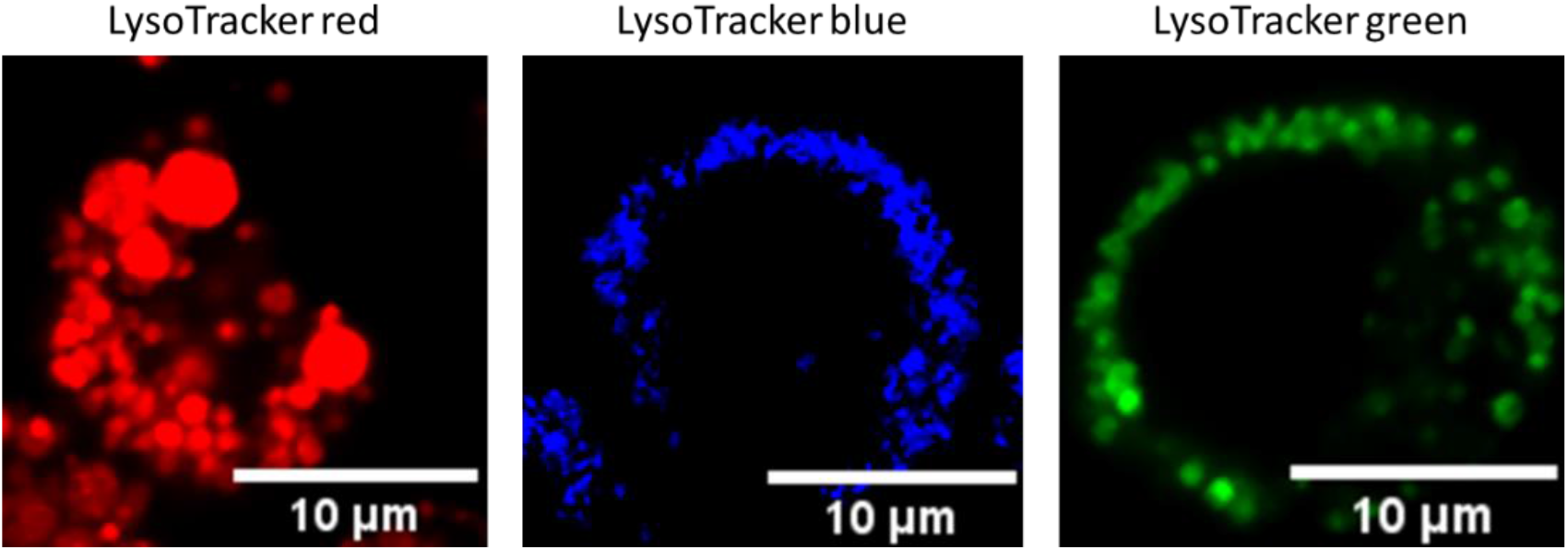
Confocal microscopy images representing different LysoTracker probes.

**Figure S3.**
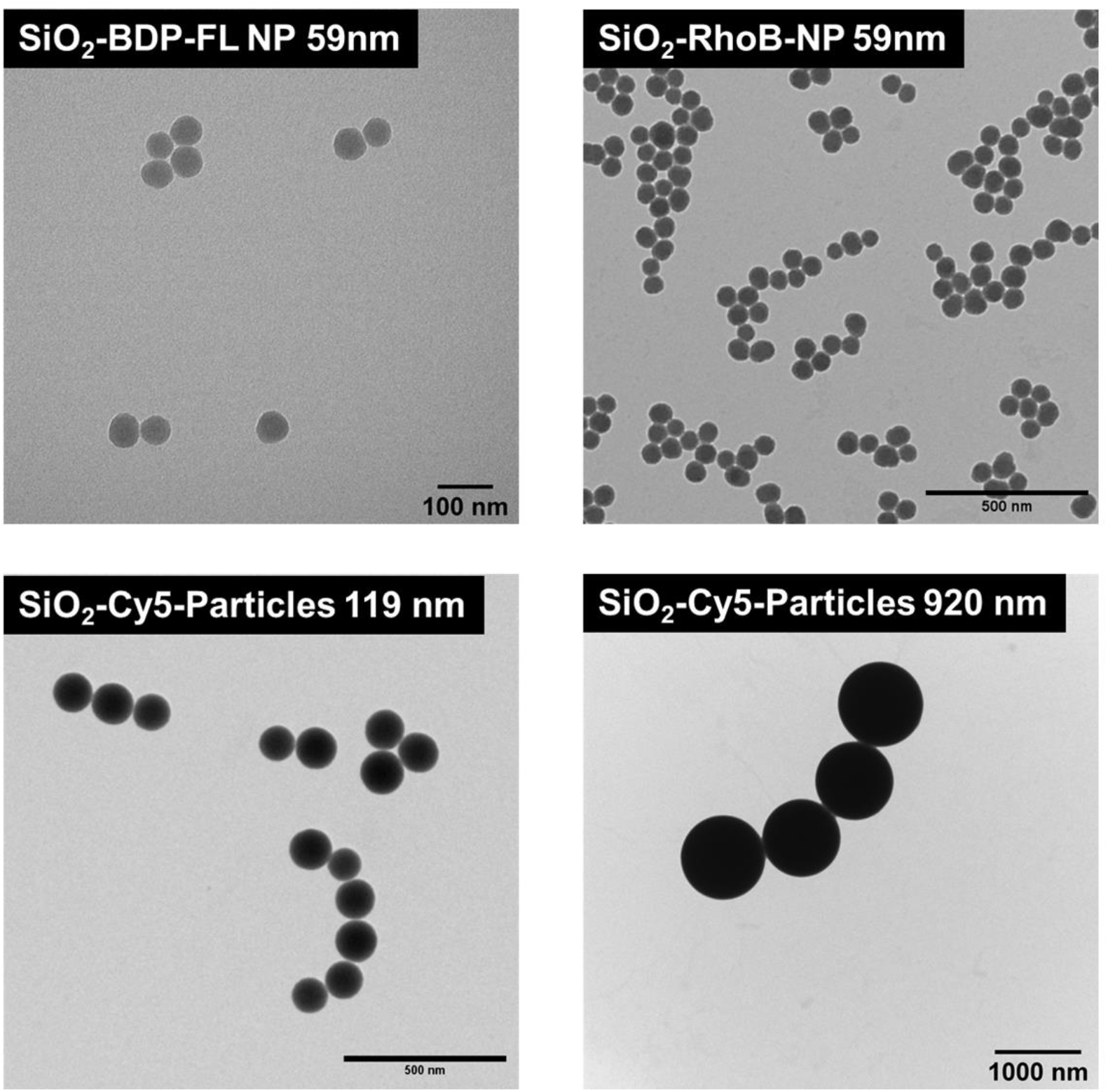
Transmission electron micrographs (TEM) of the different silica (nano)particles: 59 nm SiO_2_-BDP-FL NP, 59 nm SiO_2_-RhoB NP, 119 nm SiO_2_-Cy5 particles and 920 nm SiO_2_-Cy5 particles.

**Figure S4A.**
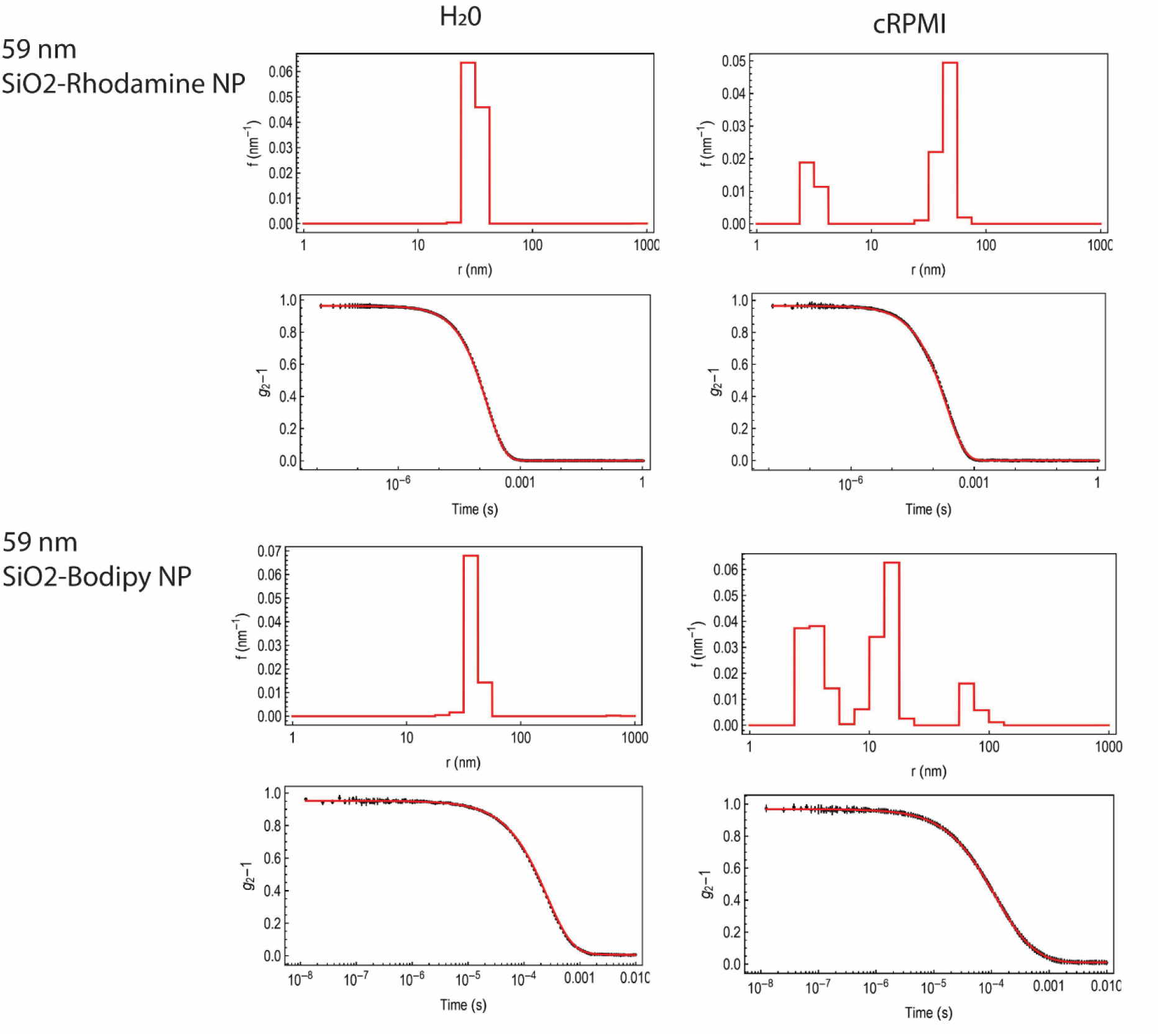
Size distributions of 59 nm SiO_2_-BDP FL NP and 59 nm SiO_2_-RhoB NP measured by dynamic light scattering in Milli-Q water and cRPMI. Representative autocorrelation functions for the DLS results are included below each histogram.

**Figure S4B.**
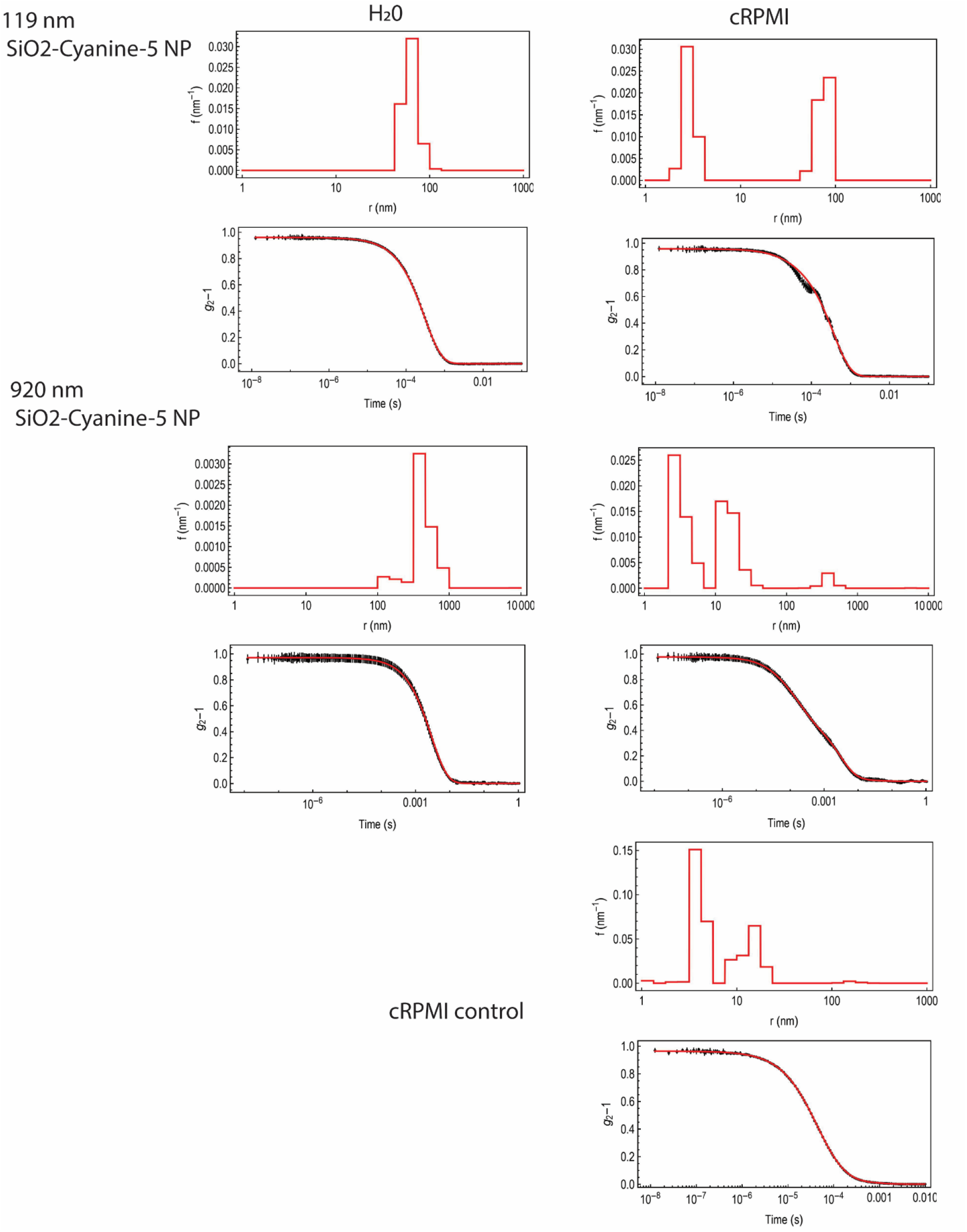
Size distributions of 119 nm SiO_2_-Cy5 particles and 920 nm SiO_2_-Cy5 particles measured by dynamic light scattering in Milli-Q water and cRPMI. Representative autocorrelation functions for the DLS results are included below each histogram.

**Figure S5A.**
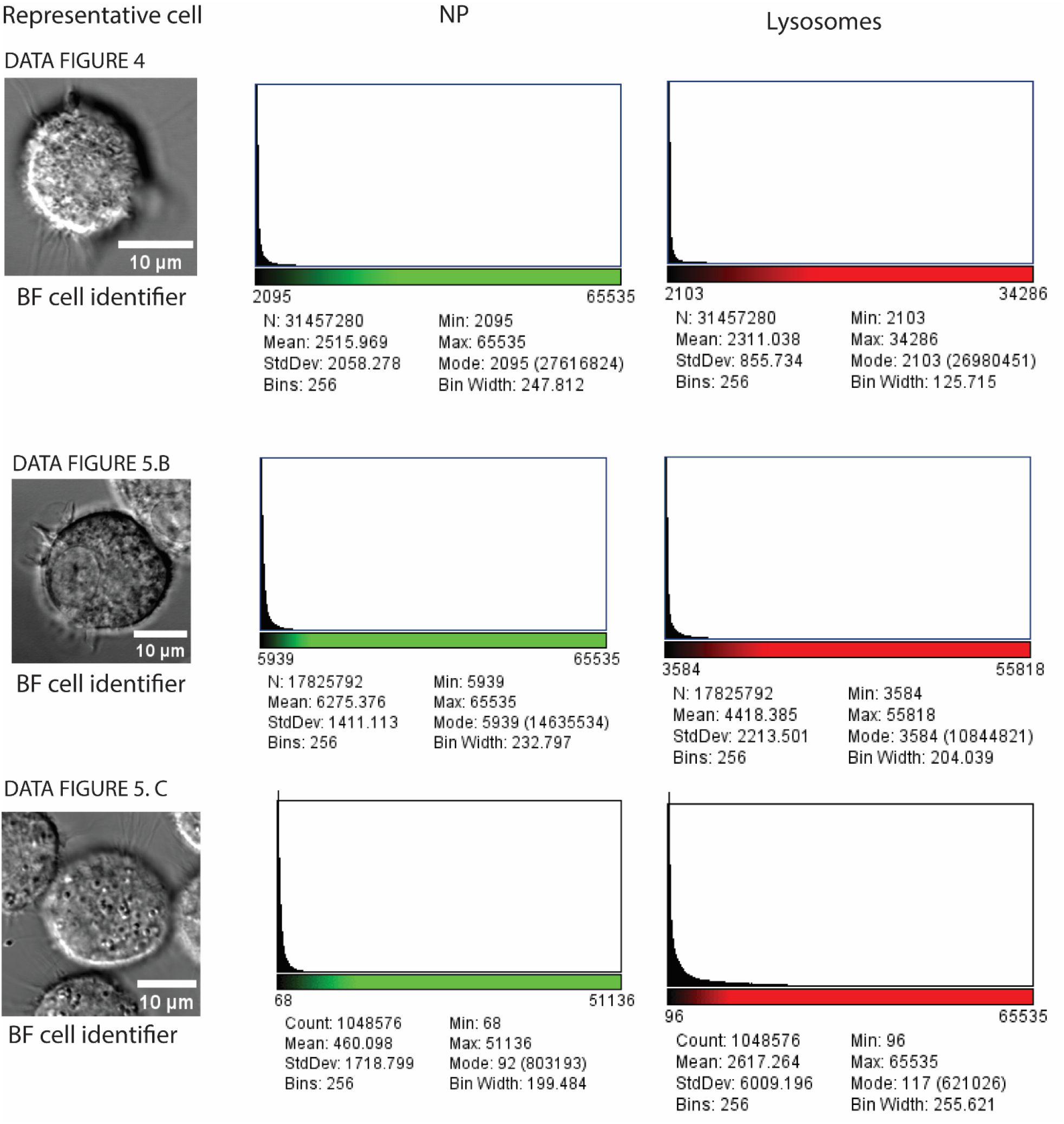
Representative images of individual cells from different experiments for identification and contour. The corresponding histograms of the complete image acquired for each individual channel (nanoparticles and lysosomes) are shown to confirm that there was no pixel overexposure. BF: Bright field image.

**Figure S5B.**
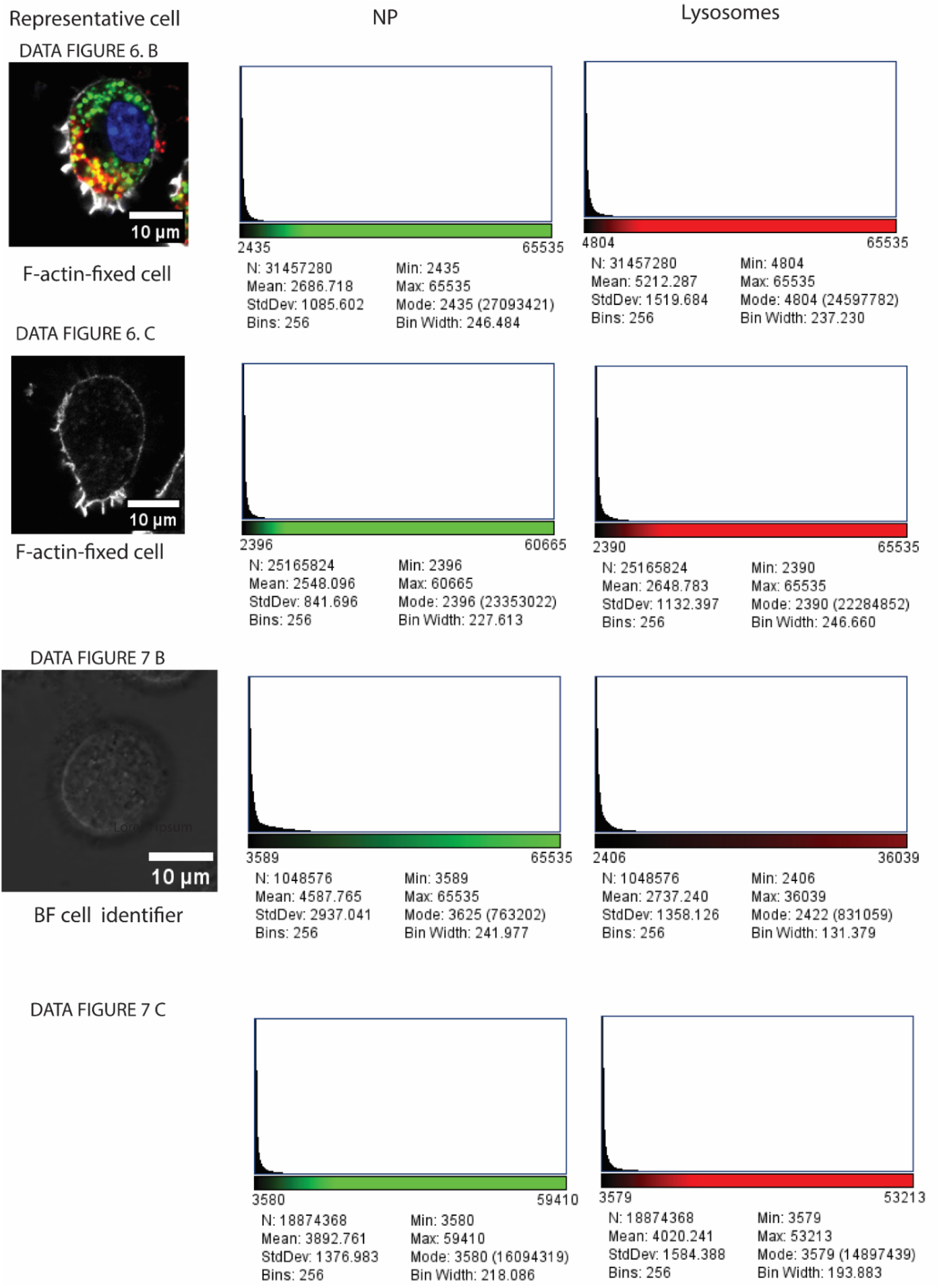
Representative images of individual cells for identification and contour with the corresponding histogram of the complete image acquired for each individual channel (nanoparticles and lysosomes). BF: Bright field image, F-actin (shown in white).

**Figure S6.**
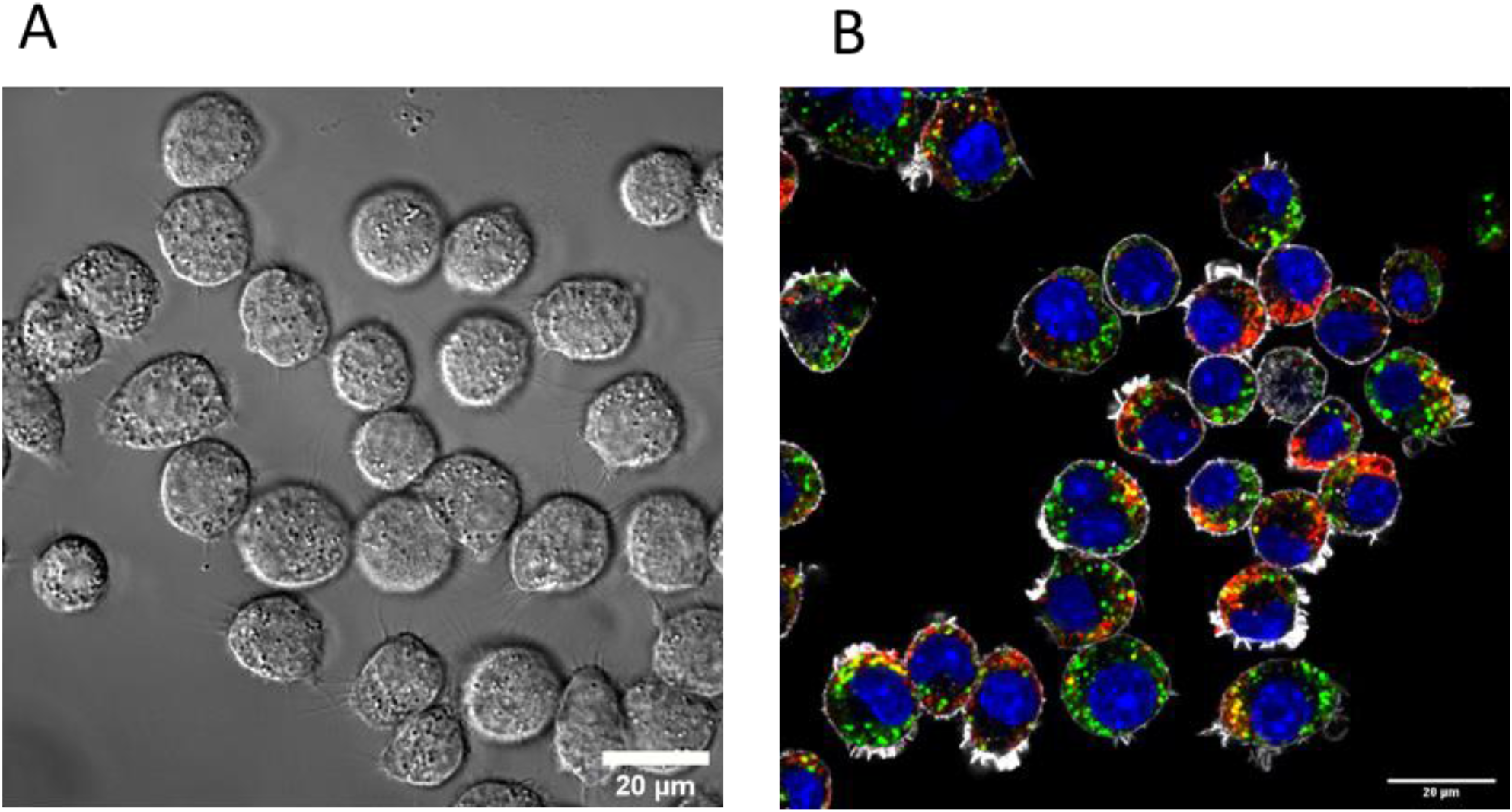
Representative bright field image of live cells **(A)** and fluorescent microscopy image of fixed cells **(B)** showing a population of individual cells used for colocalization analysis. Fixed cells were stained to identify cytoskeleton (F-actin; white), nuclei (DAPI; blue), NP (green) and lysosomes (LAMP-2, red).

**Figure S7.**
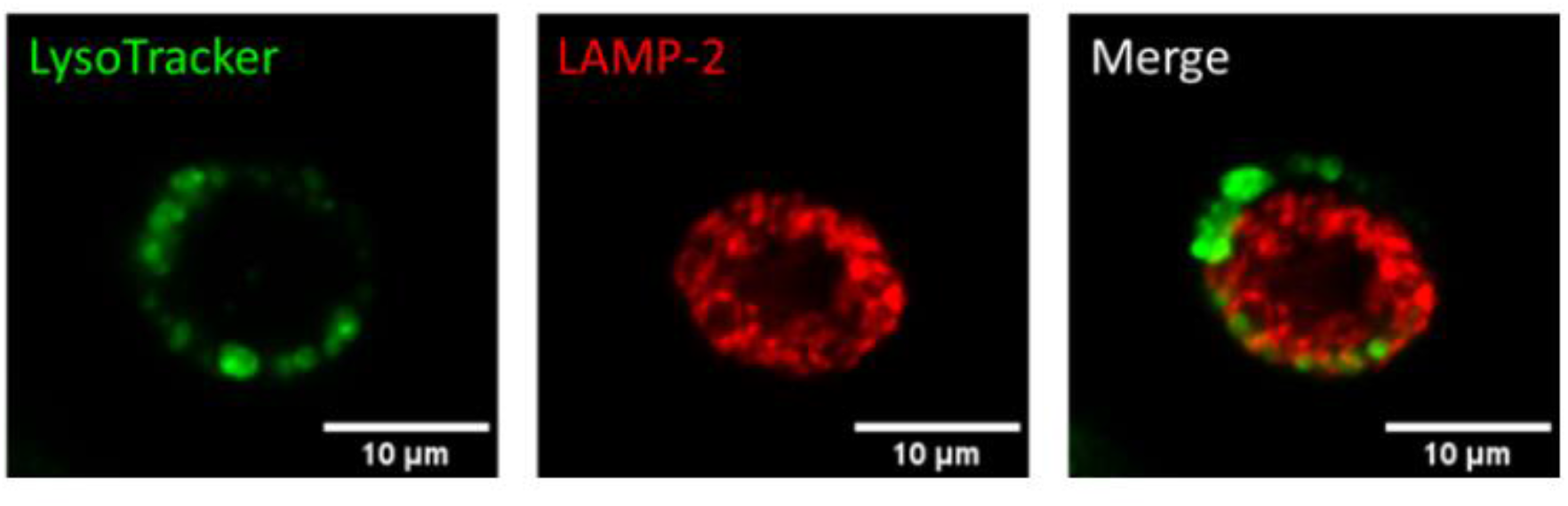
Representative images of LysoTracker Red probe and Lamp-2 stainings in a single cell. Left: Lysosomes stained with LysoTracker Red probe (green), middle: Lysosomes stained with Lamp-2 antibody (red) and right: merged channels of LysoTracker Red probe and Lamp-2 of the same cell.

**Figure S8.**
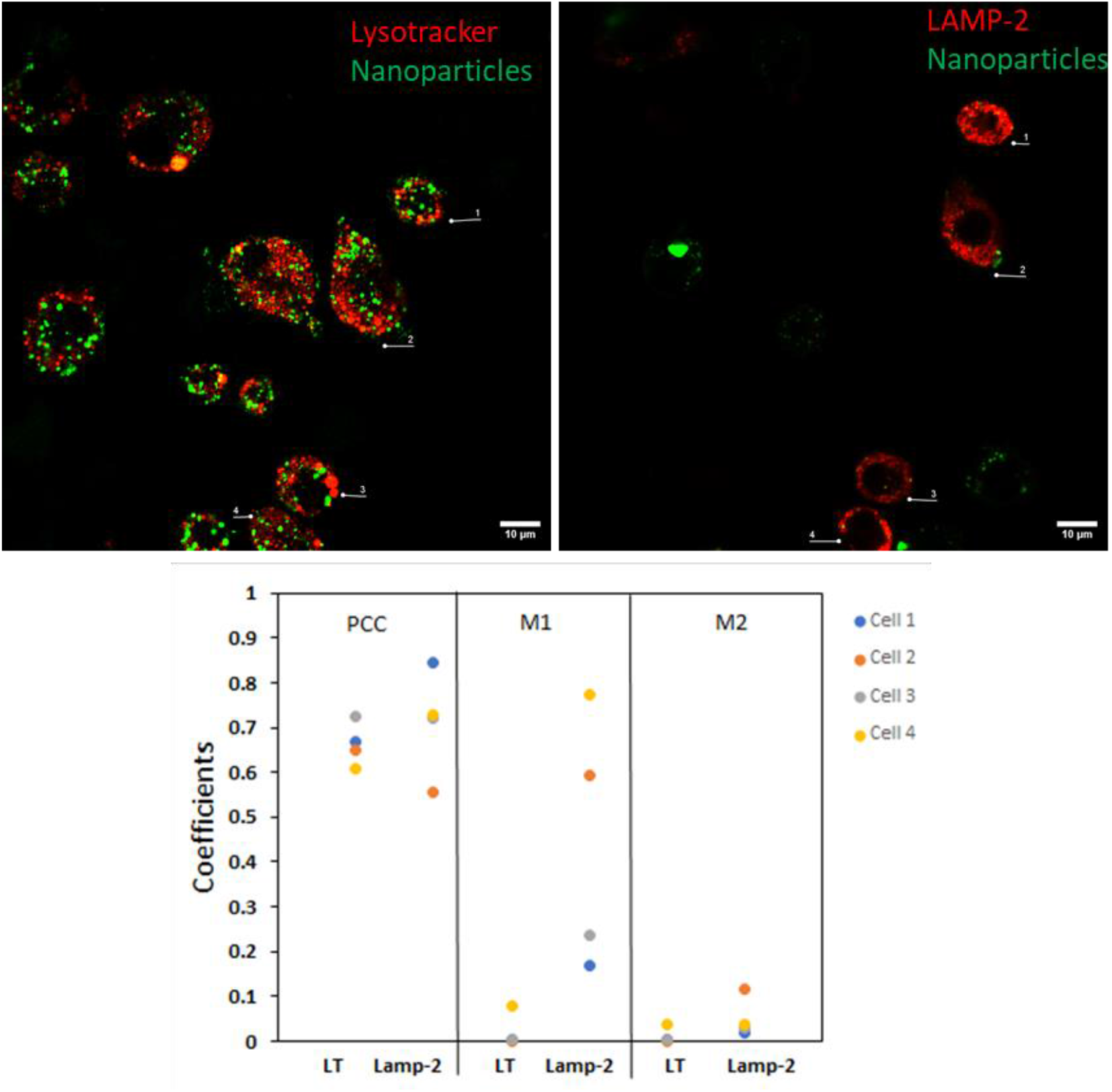
Comparison of Pearson’s and Manders’ coefficients between live and fixed cells. The cells were exposed to 59 nm SiO_2_ –BDP FL NP for 1h, stained with LysoTracker Red probe and imaged by CLSM. The same sample was fixed, stained for LAMP-2 and imaged. A total of 4 cells were analyzed for comparision, each cell has a representative color as it is shown in the coefficients plot. Each cell varies its colocalization coefficient for the NP with the lysosomes. This data demonstrates that the coefficients from live and fixed cells cannot be compared. Additionally, Pearson correlation coefficients is not enough to confirm the distribution of the fraction of the NP between the lysosomes.

**Figure S9.**
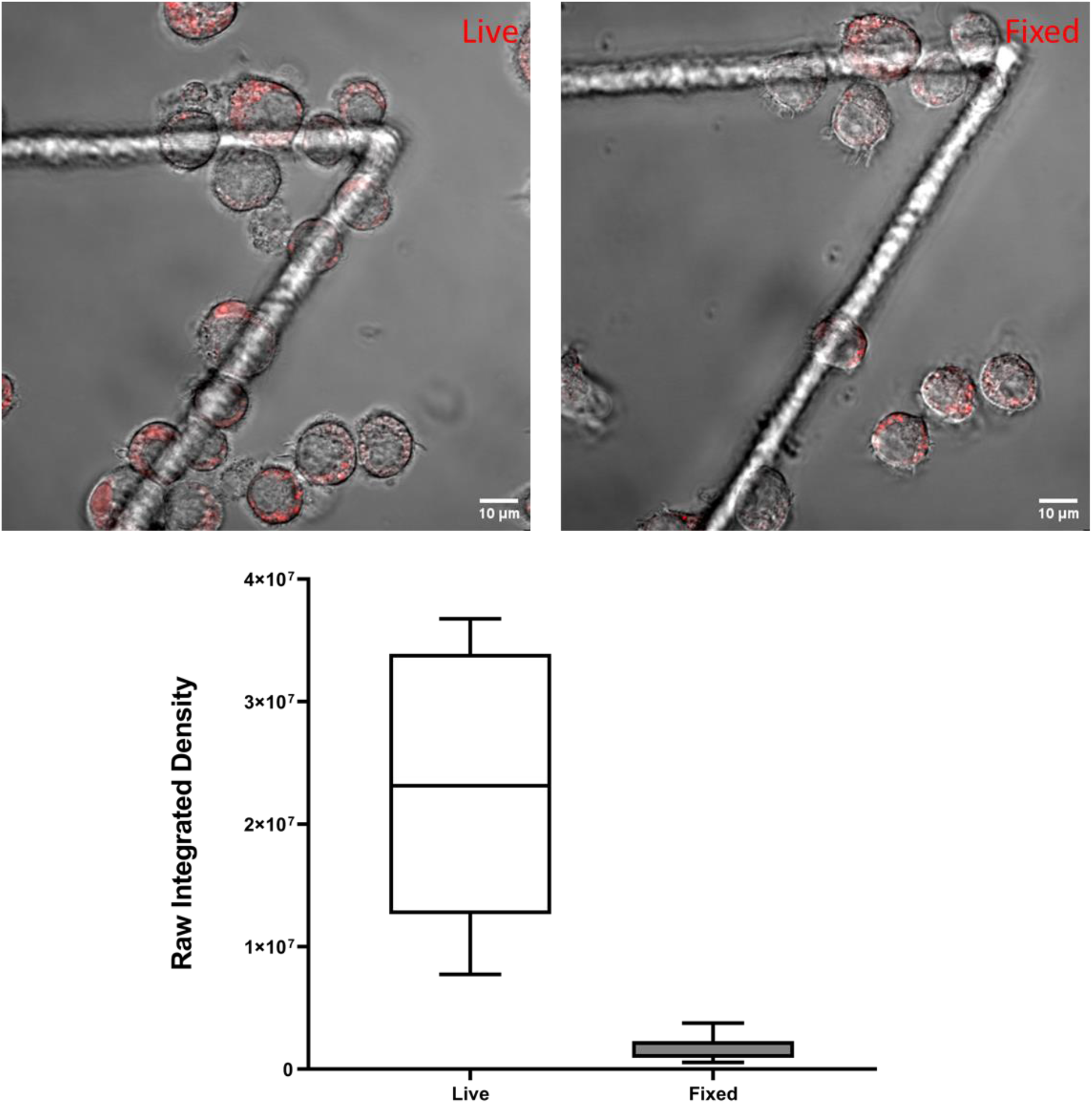
Comparison of raw integrated intensity between live and fixed cells. Top two images represent CLSM data from live cells (left) and fixed cells (right). The cells were stained with LysoTracker Red probe for 1h and imaged. The same cells were fixed and washed following a fixation protocol and imaged using the same settings. At the bottom the boxplot shows the raw integrated densities of live and fixed cells. The whiskers represent the standard deviations. Analysis was performed in 8 individual cells for each experiment. All the images were analyzed using ImageJ. There is a significant decrease in LysoTracker Red intensity between live and fixed cells.

**Video S1.**
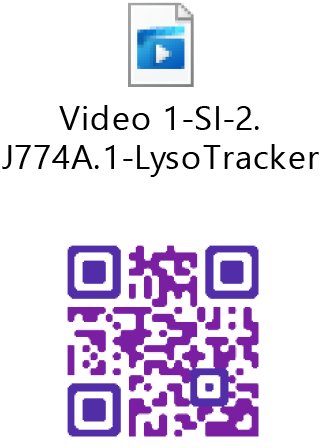
Continuous live cell imaging of J774A.1 cells, stained with LysoTracker Red for 24 h, showing increased cell death over time.

### **Script S1**. Raw integrated densities

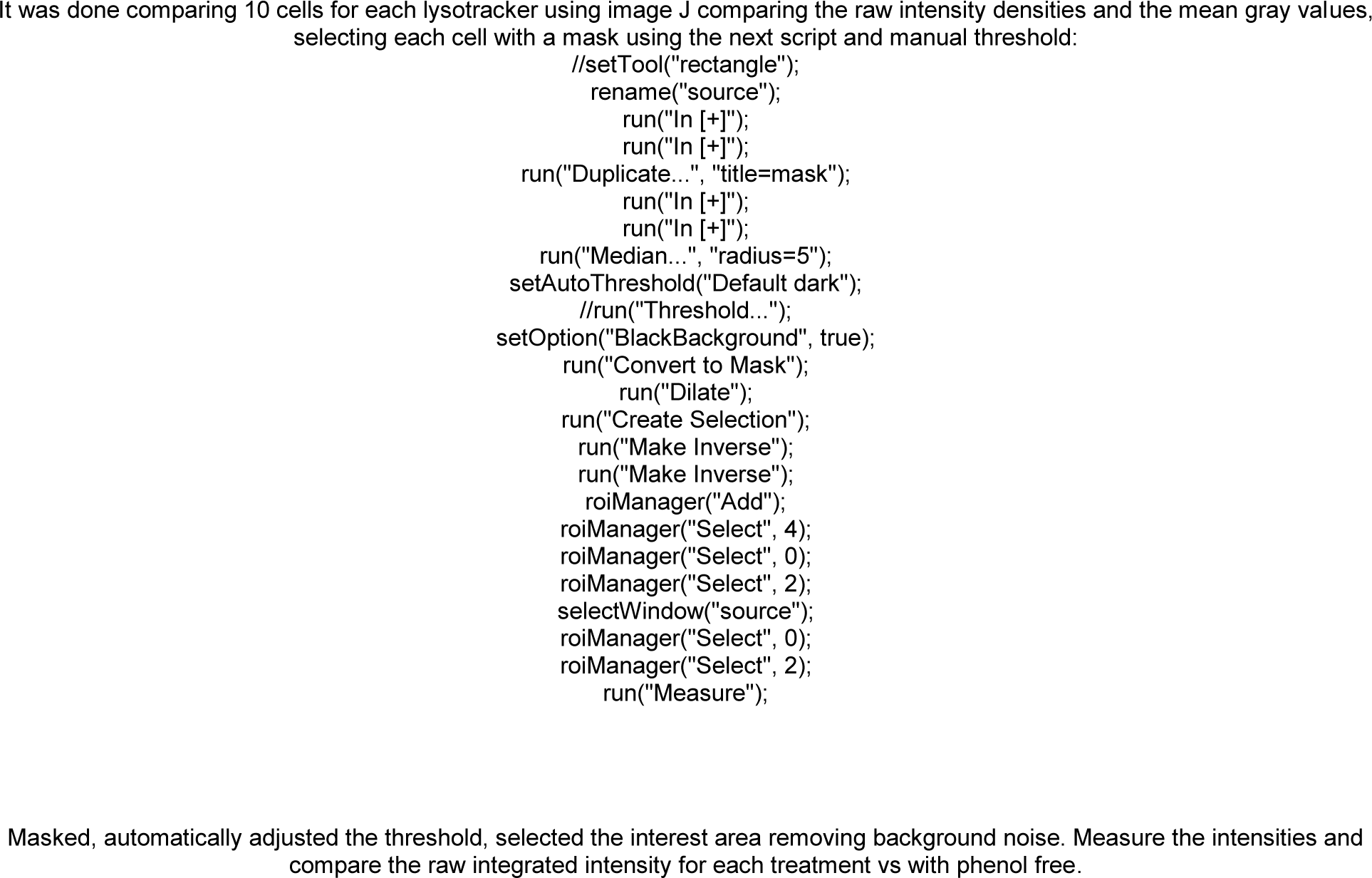

### **Script S2**. Colocalization analysis

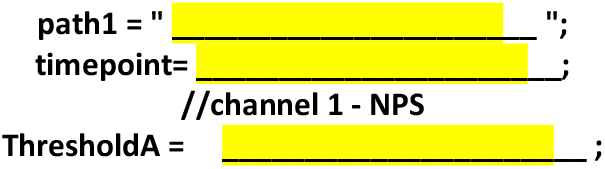
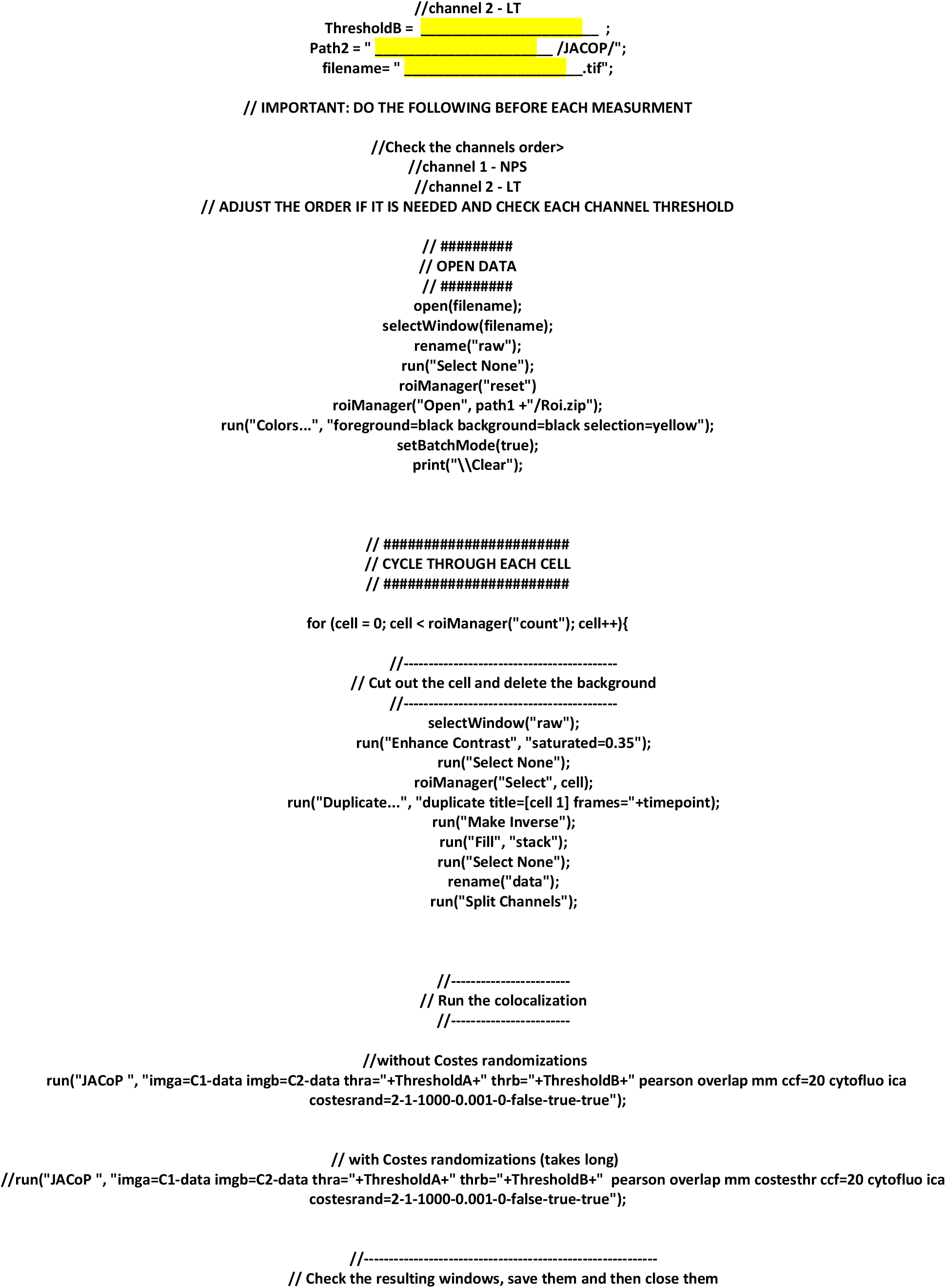
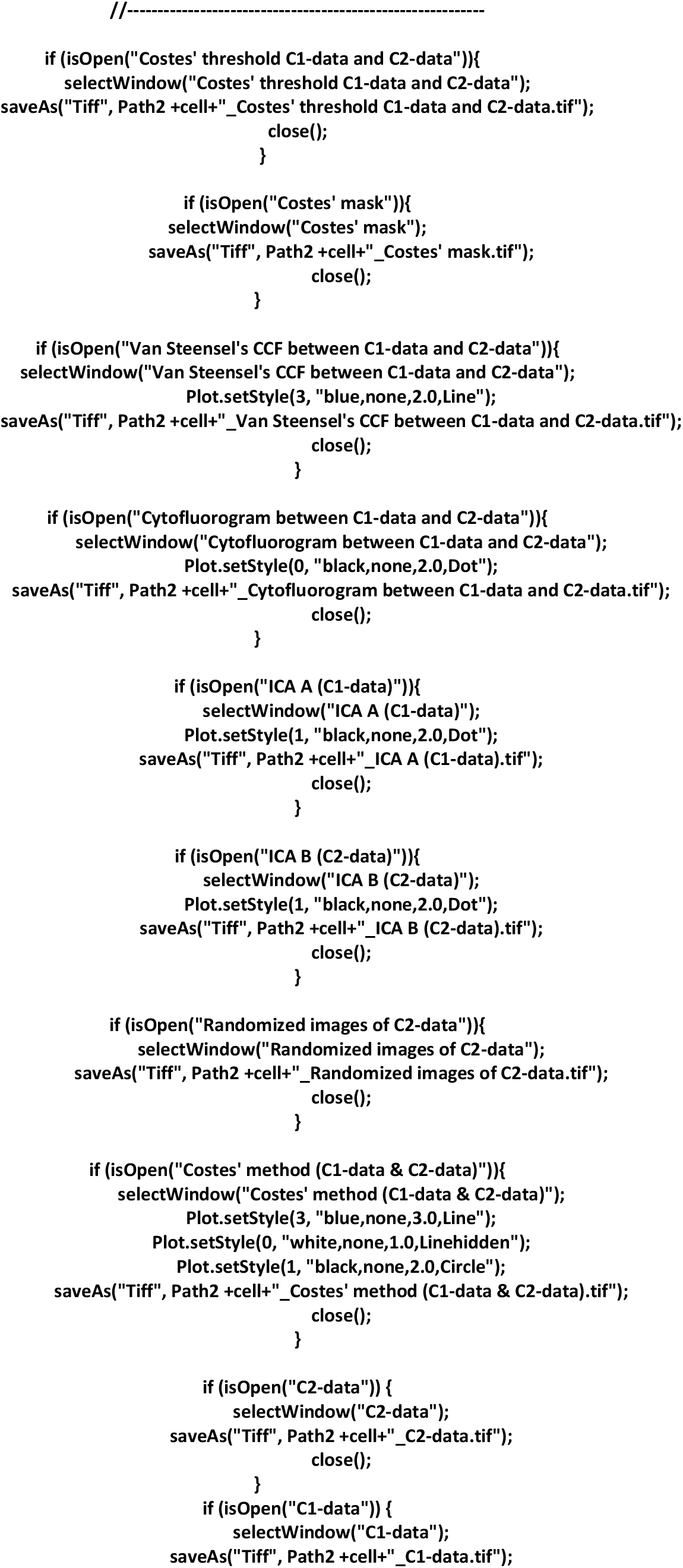
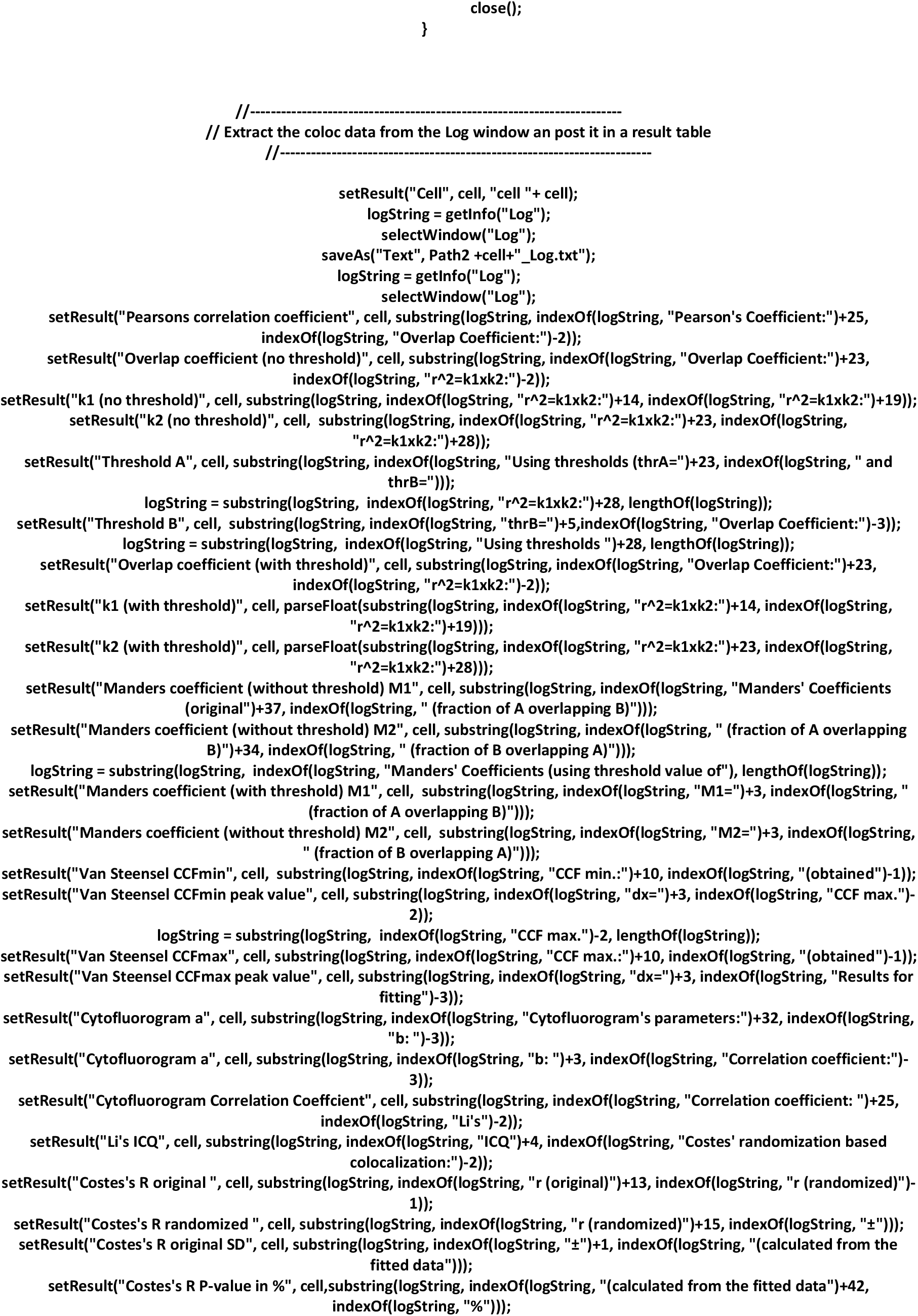
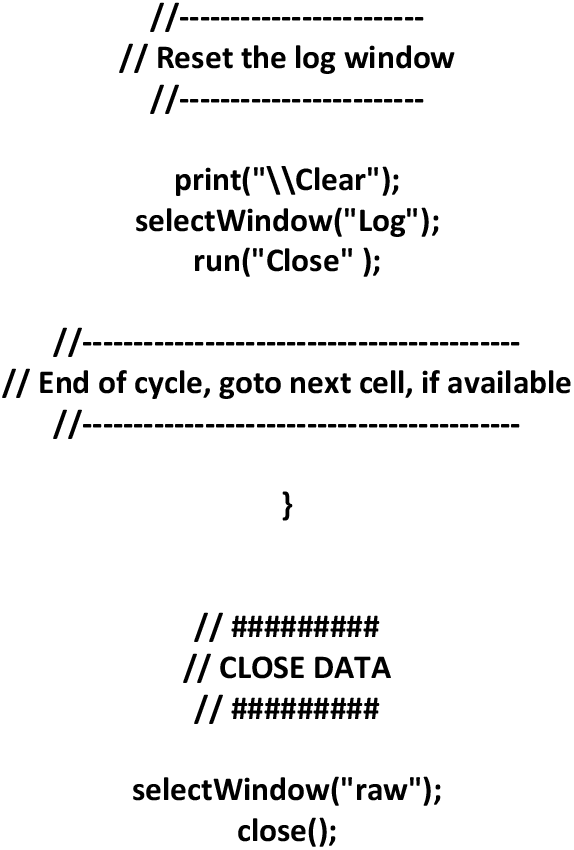

## References

1. Zhao J, Stenzel MH. Entry of nanoparticles into cells: The importance of nanoparticle properties. Polym Chem. 2018;9(3):259–272. doi:10.1039/c7py01603d

2. Park JH, Oh N. Endocytosis and exocytosis of nanoparticles in mammalian cells. Int J Nanomedicine. Published online 2014:51. doi:10.2147/ijn.s26592

3. Iversen TG, Skotland T, Sandvig K. Endocytosis and intracellular transport of nanoparticles: Present knowledge and need for future studies. Nano Today. 2011;6(2):176–185. doi:10.1016/j.nantod.2011.02.003

4. Almeida MS De, Susnik E, Drasler B, Rothen-rutishauser B. Understanding nanoparticle endocytosis to improve targeting strategies in nanomedicine. Published online 2021. doi:10.1039/d0cs01127d

5. Rennick JJ, Johnston APR, Parton RG. Key principles and methods for studying the endocytosis of biological and nanoparticle therapeutics. Nat Nanotechnol. 2021;16(3):266–276. doi:10.1038/s41565-021-00858-8

6. Brandenberger C, Mühlfeld C, Ali Z, et al. Quantitative Evaluation of Cellular Uptake and Trafficking of Plain and Polyethylene Glycol-Coated Gold Nanoparticles. Small. 2010;6(15):1669–1678. doi:10.1002/smll.201000528

7. Treuel L, Jiang X, Nienhaus GU. New views on cellular uptake and trafficking of manufactured nanoparticles. J R Soc Interface. 2013;10(82):20120939. doi:10.1098/rsif.2012.0939

8. Cupic KI, Rennick JJ, Johnston AP, Such GK. Controlling endosomal escape using nanoparticle composition: current progress and future perspectives. Nanomedicine. 2019;14(2):215–223. doi:10.2217/nnm-2018-0326

9. Saftig P, Klumperman J. Lysosome biogenesis and lysosomal membrane proteins: trafficking meets function. Nat Rev Mol Cell Biol. 2009;10(9):623–635. doi:10.1038/nrm2745

10. Ding L, Zhu X, Wang Y, et al. Intracellular Fate of Nanoparticles with Polydopamine Surface Engineering and a Novel Strategy for Exocytosis-Inhibiting, Lysosome Impairment-Based Cancer Therapy. Nano Lett. 2017;17(11):6790–6801. doi:10.1021/acs.nanolett.7b03021

11. Wong C-O, Gregory S, Hu H, et al. Lysosomal Degradation Is Required for Sustained Phagocytosis of Bacteria by Macrophages. Cell Host Microbe. 2017;21(6):719-730.e6. doi:10.1016/j.chom.2017.05.002

12. Platt FM. Sphingolipid lysosomal storage disorders. Nature. 2014;510(7503):68–75. doi:10.1038/nature13476

13. Kim SE, Overholtzer M. Autophagy proteins regulate cell engulfment mechanisms that participate in cancer. Semin Cancer Biol. 2013;23(5):329–336. doi:10.1016/j.semcancer.2013.05.004

14. Jiang P, Mizushima N. Autophagy and human diseases. Cell Res. 2014;24(1):69–79. doi:10.1038/cr.2013.161

15. Appelqvist H, Wäster P, Kågedal K, Öllinger K. The lysosome: from waste bag to potential therapeutic target. J Mol Cell Biol. 2013;5(4):214–226. doi:10.1093/jmcb/mjt022

16. Padh H, Niraj Sakhrani. Organelle targeting: third level of drug targeting. Drug Des Devel Ther. Published online July 2013:585. doi:10.2147/DDDT.S45614

17. Drasler B, Vanhecke D, Rodriguez-Lorenzo L, Petri-Fink A, Rothen-Rutishauser B. Quantifying nanoparticle cellular uptake: which method is best? Nanomedicine. 2017;12(10):1095–1099. doi:10.2217/nnm-2017-0071

18. Brandenberger C, Mühlfeld C, Ali Z, et al. Quantitative Evaluation of Cellular Uptake and Trafficking of Plain and Polyethylene Glycol-Coated Gold Nanoparticles. Small. 2010;6(15):1669–1678. doi:10.1002/smll.201000528

19. Calero M, Chiappi M, Lazaro-Carrillo A, et al. Characterization of interaction of magnetic nanoparticles with breast cancer cells. J Nanobiotechnology. 2015;13(1):16. doi:10.1186/s12951-015-0073-9

20. Vtyurina N, Åberg C, Salvati A. Imaging of nanoparticle uptake and kinetics of intracellular trafficking in individual cells. Nanoscale. 2021;13(23):10436–10446. doi:10.1039/d1nr00901j

21. Qiu K, Du Y, Liu J, Guan J-L, Chao H, Diao J. Super-resolution observation of lysosomal dynamics with fluorescent gold nanoparticles. Theranostics. 2020;10(13):6072–6081. doi:10.7150/thno.42134

22. Liu M, Li Q, Liang L, et al. Real-time visualization of clustering and intracellular transport of gold nanoparticles by correlative imaging. Nat Commun. 2017;8(1):15646. doi:10.1038/ncomms15646

23. Guehrs E, Schneider M, Günther CM, et al. Quantification of silver nanoparticle uptake and distribution within individual human macrophages by FIB/SEM slice and view. J Nanobiotechnology. 2017;15(1):21. doi:10.1186/s12951-017-0255-8

24. Tharkeshwar AK, Demedts D, Annaert W. Superparamagnetic Nanoparticles for Lysosome Isolation to Identify Spatial Alterations in Lysosomal Protein and Lipid Composition. STAR Protoc. 2020;1(3):100122. doi:10.1016/j.xpro.2020.100122

25. Zhu M, Nie G, Meng H, Xia T, Nel A, Zhao Y. Physicochemical Properties Determine Nanomaterial Cellular Uptake, Transport, and Fate. Acc Chem Res. 2013;46(3):622–631. doi:10.1021/ar300031y

26. Milosevic AM, Rodriguez-Lorenzo L, Balog S, Monnier CA, Petri-Fink A, Rothen-Rutishauser B. Assessing the Stability of Fluorescently Encoded Nanoparticles in Lysosomes by Using Complementary Methods. Angew Chemie Int Ed. 2017;56(43):13382–13386. doi:10.1002/anie.201705422

27. Chen X, Bi Y, Wang T, et al. Lysosomal Targeting with Stable and Sensitive Fluorescent Probes (Superior LysoProbes): Applications for Lysosome Labeling and Tracking during Apoptosis. Sci Rep. 2015;5:1–10. doi:10.1038/srep09004

28. Baba K, Kuwada S, Nakao A, et al. Different localization of lysosomal-associated membrane protein 1 (LAMP1) in mammalian cultured cell lines. Histochem Cell Biol. 2020;153(4):199–213. doi:10.1007/s00418-019-01842-z

29. Arruda LB, Sim D, Chikhlikar PR, et al. Dendritic Cell-Lysosomal-Associated Membrane Protein (LAMP) and LAMP-1-HIV-1 Gag Chimeras Have Distinct Cellular Trafficking Pathways and Prime T and B Cell Responses to a Diverse Repertoire of Epitopes. J Immunol. 2006;177(4):2265–2275. doi:10.4049/jimmunol.177.4.2265

30. Bourquin J, Septiadi D, Vanhecke D, et al. Reduction of Nanoparticle Load in Cells by Mitosis but Not Exocytosis. ACS Nano. 2019;13(7):7759–7770. doi:10.1021/acsnano.9b01604

31. Schütz I, Lopez-Hernandez T, Gao Q, et al. Lysosomal dysfunction caused by cellular accumulation of silica nanoparticles. J Biol Chem. 2016;291(27):14170–14184. doi:10.1074/jbc.M115.710947

32. Susnik E, Taladriz-Blanco P, Drasler B, Balog S, Petri-Fink A, Rothen-Rutishauser B. Increased Uptake of Silica Nanoparticles in Inflamed Macrophages but Not upon Co-Exposure to Micron-Sized Particles. Cells. 2020;9(9). doi:10.3390/cells9092099

33. Sugimoto Y, Ninomiya H, Ohsaki Y, et al. Accumulation of cholera toxin and GM1 ganglioside in the early endosome of niemann-pick C1-deficient cells. Proc Natl Acad Sci U S A. 2001;98(22):12391–12396. doi:10.1073/pnas.221181998

34. Freundt EC, Czapiga M, Lenardo MJ. Photoconversion of Lysotracker Red to a green fluorescent molecule. Cell Res. 2007;17(11):956–958. doi:10.1038/cr.2007.80

35. Kim D-H, Chang Y, Park S, et al. Blue-conversion of organic dyes produces artifacts in multicolor fluorescence imaging. Chem Sci. 2021;12(25):8660–8667. doi:10.1039/d1sc00612f

36. Aaron JS, Taylor AB, Chew T-L. Image co-localization – co-occurrence versus correlation. J Cell Sci. 2018;131(3):jcs211847. doi:10.1242/jcs.211847

37. Manders EMM, Verbeek FJ, Aten JA. Measurement of co-localization of objects in dual-colour confocal images. J Microsc. 1993;169(3):375–382. doi:10.1111/j.1365-2818.1993.tb03313.x

38. Pearson K. VII. Mathematical contributions to the theory of evolution.—III. Regression, heredity, and panmixia. Philos Trans R Soc London Ser A, Contain Pap a Math or Phys Character. 1896;187:253–318. doi:10.1098/rsta.1896.0007

39. Adler J, Pagakis SN, Parmryd I. Replicate-based noise corrected correlation for accurate measurements of colocalization. J Microsc. 2008;230(1):121–133. doi:10.1111/j.1365-2818.2008.01967.x

40. Costes S V., Daelemans D, Cho EH, Dobbin Z, Pavlakis G, Lockett S. Automatic and Quantitative Measurement of Protein-Protein Colocalization in Live Cells. Biophys J. 2004;86(6):3993–4003. doi:10.1529/biophysj.103.038422

41. Li Q. A Syntaxin 1, G o, and N-Type Calcium Channel Complex at a Presynaptic Nerve Terminal: Analysis by Quantitative Immunocolocalization. J Neurosci. 2004;24(16):4070–4081. doi:10.1523/JNEUROSCI.0346-04.2004

42. Woodcroft BJ, Hammond L, Stow JL, Hamilton NA. Automated organelle-based colocalization in whole-cell imaging. Cytom Part A. 2009;75A(11):941-950. doi:10.1002/cyto.a.20786

43. Kuhn DA, Vanhecke D, Michen B, et al. Different endocytotic uptake mechanisms for nanoparticles in epithelial cells and macrophages. Beilstein J Nanotechnol. 2014;5(1):1625–1636. doi:10.3762/bjnano.5.174

44. Gallud A, Bondarenko O, Feliu N, et al. Macrophage activation status determines the internalization of mesoporous silica particles of different sizes: Exploring the role of different pattern recognition receptors. Biomaterials. 2017;121:28–40. doi:10.1016/j.biomaterials.2016.12.029

45. Schrijvers DM, Martinet W, De Meyer GRY, Andries L, Herman AG, Kockx MM. Flow cytometric evaluation of a model for phagocytosis of cells undergoing apoptosis. J Immunol Methods. 2004;287(1-2):101-108. doi:10.1016/j.jim.2004.01.013

46. Clift MJD, Rothen-Rutishauser B, Brown DM, et al. The impact of different nanoparticle surface chemistry and size on uptake and toxicity in a murine macrophage cell line. Toxicol Appl Pharmacol. 2008;232(3):418–427. doi:10.1016/j.taap.2008.06.009

47. Scientific TF. Fluorescence SpectraViewer. Published 2022. https://www.thermofisher.com/order/fluorescence-spectraviewer#!/

48. Ashino Y, Ying X, Dobbs LG, Bhattacharya J. [Ca 2+] i oscillations regulate type II cell exocytosis in the pulmonary alveolus. Am J Physiol Cell Mol Physiol. 2000;279(1):L5–L13. doi:10.1152/ajplung.2000.279.1.L5

49. Wemhöner A, Frick M, Dietl P, Jennings P, Haller T. A Fluorescent Microplate Assay for Exocytosis in Alveolar Type II Cells. J Biomol Screen. 2006;11(3):286–295. doi:10.1177/1087057105285284

50. Li Y, Wang S, Zhao Y, et al. A Model In Vitro Study Using Hypericin: Tumor-Versus Necrosis-Targeting Property and Possible Mechanisms. Biology (Basel). 2020;9(1):13. doi:10.3390/biology9010013

51. Viswanathan G, Hsu Y-H, Voon SH, et al. A Comparative Study of Cellular Uptake and Subcellular Localization of Doxorubicin Loaded in Self-Assemblies of Amphiphilic Copolymers with Pendant Dendron by MDA-MB-231 Human Breast Cancer Cells. Macromol Biosci. 2016;16(6):882–895. doi:10.1002/mabi.201500435

52. Bussi C, Peralta Ramos JM, Arroyo DS, et al. Alpha-synuclein fibrils recruit TBK1 and OPTN to lysosomal damage sites and induce autophagy in microglial cells. J Cell Sci. Published online January 1, 2018. doi:10.1242/jcs.226241

53. Xu H, Lee S-J, Suzuki E, et al. A lysosomal tetraspanin associated with retinal degeneration identified via a genome-wide screen. EMBO J. 2004;23(4):811–822. doi:10.1038/sj.emboj.7600112

54. Pol A, Luetterforst R, Lindsay M, Heino S, Ikonen E, Parton RG. A Caveolin Dominant Negative Mutant Associates with Lipid Bodies and Induces Intracellular Cholesterol Imbalance. J Cell Biol. 2001;152(5):1057–1070. doi:10.1083/jcb.152.5.1057

55. Lou X, Zhang M, Zhao Z, et al. A photostable AIE fluorogen for lysosome-targetable imaging of living cells. J Mater Chem B. 2016;4(32):5412–5417. doi:10.1039/c6tb01293k

56. Kinnear C, Moore TL, Rodriguez-Lorenzo L, Rothen-Rutishauser B, Petri-Fink A. Form Follows Function: Nanoparticle Shape and Its Implications for Nanomedicine. Chem Rev. 2017;117(17):11476–11521. doi:10.1021/acs.chemrev.7b00194

57. Jonkman J, Brown CM, Wright GD, Anderson KI, North AJ. Tutorial: guidance for quantitative confocal microscopy. Nat Protoc. 2020;15(5):1585–1611. doi:10.1038/s41596-020-0313-9

58. Russ JC. The Image Processing Handbook. 6th ed.; 2016.

59. Leong FJW-M. Correction of uneven illumination (vignetting) in digital microscopy images. J Clin Pathol. 2003;56(8):619–621. doi:10.1136/jcp.56.8.619

60. Leng S, Bruesewitz M, Tao S, et al. Photon-counting Detector CT: System Design and Clinical Applications of an Emerging Technology. RadioGraphics. 2019;39(3):729–743. doi:10.1148/rg.2019180115

61. Waters JC. Accuracy and precision in quantitative fluorescence microscopy. J Cell Biol. 2009;185(7):1135–1148. doi:10.1083/jcb.200903097

62. Chikte S, Panchal N, Warnes G. Use of LysoTracker dyes: A flow cytometric study of autophagy. Cytom Part A. 2014;85(2):169–178. doi:10.1002/cyto.a.22312

63. Kim JA, Aberg C, Salvati A, Dawson KA. Role of cell cycle on the cellular uptake and dilution of nanoparticles in a cell population. Nat Nanotechnol. 2012;7(1):62–68. doi:10.1038/nnano.2011.191

64. Åberg C. Kinetics of nanoparticle uptake into and distribution in human cells. Nanoscale Adv. 2021;3(8):2196–2212. doi:10.1039/d0na00716a

65. Cheng W, Kim S, Zivkovic S, Chung H, Ren Y, Guan J. Specific Labelling of Phagosome-Derived Vesicles in Macrophages with a Membrane Dye Delivered with Microfabricated Microparticles. SSRN Electron J. Published online 2021:1-30. doi:10.2139/ssrn.3937057

66. Li Y, Hu Q, Miao G, et al. Size-dependent mechanism of intracellular localization and cytotoxicity of mono-disperse spherical mesoporous nano-and micron-bioactive glass particles. J Biomed Nanotechnol. 2016;12(5):863–877. doi:10.1166/jbn.2016.2235

67. Seydoux E, Rothen-Ruthishauser B, Nita I, et al. Size-dependent accumulation of particles in lysosomes modulates dendritic cell function through impaired antigen degradation. Int J Nanomedicine. Published online August 2014:3885. doi:10.2147/IJN.S64353

68. Oh YK, Swanson JA. Different fates of phagocytosed particles after delivery into macrophage lysosomes. J Cell Biol. 1996;132(4):585–593. doi:10.1083/jcb.132.4.585

69. Zhukova V, Osipova N, Semyonkin A, et al. Fluorescently Labeled PLGA Nanoparticles for Visualization In Vitro and In Vivo: The Importance of Dye Properties. Pharmaceutics. 2021;13(8):1145. doi:10.3390/pharmaceutics13081145

70. Rodriguez-Lorenzo L, Fytianos K, Blank F, Von Garnier C, Rothen-Rutishauser B, Petri-Fink A. Fluorescence-encoded gold nanoparticles: Library design and modulation of cellular uptake into dendritic cells. Small. 2014;10(7):1341–1350. doi:10.1002/smll.201302889

71. Hirota Y, Masuyama N, Kuronita T, Fujita H, Himeno M, Tanaka Y. Analysis of postlysosomal compartments. Biochem Biophys Res Commun. 2004;314(2):306–312. doi:10.1016/j.bbrc.2003.12.092

72. Wang F, Salvati A, Boya P. Lysosome-dependent cell death and deregulated autophagy induced by amine-modified polystyrene nanoparticles. Open Biol. 2018;4(8). doi:10.1098/rsob.170271

73. Solorio-Rodríguez A, Escamilla-Rivera V, Uribe-Ramírez M, et al. In vitro cytotoxicity study of superparamagnetic iron oxide and silica nanoparticles on pneumocyte organelles. Toxicol Vitr. 2021;72(November 2020). doi:10.1016/j.tiv.2020.105071

74. van Meel E, Bos E, van der Lienden MJC, et al. Localization of active endogenous and exogenous β-glucocerebrosidase by correlative light-electron microscopy in human fibroblasts. Traffic. 2019;20(5):346–356. doi:10.1111/tra.12641

75. Banquy X, Suarez F, Argaw A, et al. Effect of mechanical properties of hydrogel nanoparticles on macrophage cell uptake. Soft Matter. 2009;5(20):3984–3991. doi:10.1039/b821583a

76. Yu T, Malugin A, Ghandehari H. Impact of silica nanoparticle design on cellular toxicity and hemolytic activity. ACS Nano. 2011;5(7):5717–5728. doi:10.1021/nn2013904

77. Abdelkhaliq A, Van Der Zande M, Punt A, et al. Impact of nanoparticle surface functionalization on the protein corona and cellular adhesion, uptake and transport 03 Chemical Sciences 0306 Physical Chemistry (incl. Structural). J Nanobiotechnology. 2018;16(1):1–13. doi:10.1186/s12951-018-0394-6

78. Krentz AJ, Bailey CJ. Oral antidiabetic agents: Current role in type 2 diabetes mellitus. Drugs. 2005;65(3):385–411. doi:10.2165/00003495-200565030-00005

79. Ramsey N. Majzoub, Emily Wonder, Kai K. Ewert, Venkata Ramana Kotamraju, Tambet Teesalu CRS. Rab11 and Lysotracker Markers Reveal Correlation between Endosomal Pathways and Transfection Efficiency of Surface-Functionalized Cationic Liposome–DNA Nanoparticles. Physiol Behav. 2016;120(26):6439–6453. doi:10.1021/acs.jpcb.6b04441.Rab11

80. De Boer P, Hoogenboom JP, Giepmans BNG. Correlated light and electron microscopy: Ultrastructure lights up! Nat Methods. 2015;12(6):503–513. doi:10.1038/nmeth.3400

81. Moore TL, Urban DA, Rodriguez-Lorenzo L, et al. Nanoparticle administration method in cell culture alters particle-cell interaction. Sci Rep. 2019;9(1):1–9. doi:10.1038/s41598-018-36954-4

82. Bolte S, Cordelières FP. A guided tour into subcellular colocalization analysis in light microscopy. J Microsc. 2006;224(3):213–232. doi:10.1111/j.1365-2818.2006.01706.x

83. Ettinger A (2014). Fluorescence live cell imaging. Methods Cell Biol. 2014;123:77–94. doi:10.1016/B978-0-12-420138-5.00005-7

84. Stöber W, Fink A, Bohn E. Controlled growth of monodisperse silica spheres in the micron size range. J Colloid Interface Sci. 1968;26(1):62–69. doi:10.1016/0021-9797(68)90272-5

